# Cannabidiol Inhibits PIEZO Channels to Mitigate Red Blood Disorders

**DOI:** 10.64898/2026.01.22.700543

**Authors:** Pengfei Liang, Yui-Chun Serena Wan, Ke Zoe Shan, Yang Zhang, Martha Delahunty, Sanjay Khandelwal, Gowthami M. Arepally, Marilyn J. Telen, Huanghe Yang

**Author notes:** Correspondence: Huanghe Yang, Address: Box 3711, DUMC, Durham, NC 27514, USA.

## Abstract

Hyperactivity of the mechanosensitive ion channel PIEZO1 promotes pathologic Ca²⁺ overload in red blood cells (RBCs), driving dehydration, TMEM16F-dependent phosphatidylserine (PS) exposure, microparticle shedding, and increased thrombotic and vaso-occlusive risks in hereditary xerocytosis (HX) and sickle cell disease (SCD). However, clinically deployable PIEZO inhibitors to treat these blood disorders are lacking. Here we report that cannabidiol (CBD), a non-psychoactive cannabinoid commonly used in SCD patients for pain management, inhibits PIEZO1 activity and restores aberrant mechanotransduction in HX and SCD RBCs. Micromolar concentrations of CBD blocks PIEZO1 currents and suppresses PIEZO1-mediate Ca²⁺ entry. In HX and SCD RBCs, CBD attenuates PIEZO1-TMEM16F coupling, thereby reducing PS exposure, microparticle release, thrombin generation, RBC-endothelium adhesion, and sickling. Beyond RBCs, CBD also blocks PIEZO2 currents and PIEZO2-dependent mechanical sensation in mice, suggesting broader effects of CBD-mediated PIEZO inhibition on nociceptive functions. Together, our findings identify CBD as a potent PIEZO inhibitor that restores calcium and membrane homeostasis, supporting the repurposing of CBD or the development of CBD-derived, PIEZO-selective analogs as a promising disease-modifying strategy for SCD, HX, and other PIEZO-mediated mechanosensing disorders.

**Highlights:**

- CBD inhibits PIEZO channels and disrupts the PIEZO1-TMEM16F axis in diseased RBCs
- CBD shows a therapeutic window to prevent PS exposure and translational promise for HX and SCD

## INTRODUCTION

SCD remains a major global health burden, affecting over 7 million people worldwide, including about 100,000 in the United States, primarily of African and Hispanic descent^1,2^. Current therapies, including hydroxyurea, L-glutamine, blood transfusions, and newer agents like voxelotor and crizanlizumab, offer partial benefit but have considerable limitations^3,4^. Hydroxyurea and L-glutamine have less efficacy and lack mechanistic specificity for SCD. Newer targeted drugs are costly, only modestly effective across SCD complications, and associated with adverse effects. Gene and cell therapies hold promise but remain inaccessible for most patients due to their complexity, expense, and safety concerns^5,6^. These challenges underscore the urgent need for safe, affordable, and mechanistically derived treatments for SCD and related red cell disorders.

The mechanosensitive ion channel PIEZO1 has recently emerged as a key player in the pathophysiology of SCD^7–11^. In RBCs, PIEZO1 converts mechanical stress into Ca²⁺ signals that regulate cell hydration, deformability, ATP release, and membrane lipid asymmetry ^8,10,12–14^. We previously discovered that PIEZO1-mediated Ca²⁺ influx activates the Ca²⁺-dependent phospholipid scramblase TMEM16F^15,16^, resulting in loss of lipid asymmetry, exposure of PS, and microparticle release in RBCs^7,13^. Enhanced PIEZO1-TMEM16F coupling in SCD and HX, a rare RBC disorder caused by gain-of-function (GOF) mutations in PIEZO1^17–19^, drives excessive PS exposure, microvesiculation, and prothrombotic activity^7,13^. PS-exposed RBCs and microparticles provide catalytic surfaces for coagulation and promote RBC-endothelium adhesion, contributing to anemia, hypercoagulability, and vaso-occlusion. We further identified benzbromarone, an anti-gout drug, as a potent PIEZO1 inhibitor that normalizes PIEZO1-TMEM16F activity, restores Ca²⁺ and membrane homeostasis, and mitigates PS exposure in SCD and HX RBCs^7,13^. However, concerns about benzbromarone-associated hepatotoxicity limit its translational potential^20^. These findings highlight PIEZO1 as a druggable target for restoring ionic and lipid balance in diseased RBCs and motivate the search for clinically safe PIEZO1 inhibitors as a new therapeutic strategy for red cell disorders.

Here, we report our accidental finding that CBD, a non-psychoactive cannabinoid widely used for the treatment of epilepsy^21^ and pain^22,23^ including SCD-associated hyperalgesia^24^, acts as a potent inhibitor of PIEZO1 and PIEZO2. In RBCs, sub-micromolar concentrations of CBD partially inhibit PIEZO1 activity, preserving its physiological roles in maintaining RBC homeostasis while effectively disrupting PIEZO1-TMEM16F coupling and subsequent phospholipid scrambling. As a result, CBD markedly reduces PS exposure, microparticle release, thrombin generation, RBC-endothelium adhesion, hemolysis, and sickling, in HX and SCD RBCs. Beyond RBCs, CBD also blocks PIEZO2 and inhibits PIEZO2-dependent mechanosensation in mice, suggesting broader implications for CBD-mediated PIEZO inhibition in modulating nociceptive functions. Together, these findings identify CBD as a naturally occurring PIEZO inhibitor that restores calcium and membrane homeostasis, supporting the repurposing of CBD or development of CBD-derived PIEZO-selective analogs as a promising, disease-modifying therapeutic strategy for SCD, HX, and other PIEZO-mediated mechanosensing disorders.

## RESULTS

### CBD abolishes PIEZO1-mediated Ca^2+^ entry and subsequent PS exposure in RBCs

Our initial motivation to test CBD, a known agonist for TRPV2^25^, was to examine whether the Ca^2+^ entry through this newly identified cation channel in RBCs^26,27^ could act in concert with PIEZO1-mediated Ca^2+^ entry to synergistically promote TMEM16F-dependent lipid scrambling and PS exposure (Fig. 1A). Contrary to our hypothesis that co-activation of PIEZO1 and TRPV2 by their respective agonists, Yoda1 and CBD, would enhance RBC dehydration and PS exposure, we found that 200 µM CBD completely abolished 2 µM Yoda1-induced Ca^2+^ entry (Fig. S1) and the subsequent PS exposure in healthy donor (HD) RBCs, as measured both at a population level using a plate reader and at single-cell resolution by fluorescence microscopy^13^ (Figs. 1B-E and Video S1-2).

**Figure 1.**
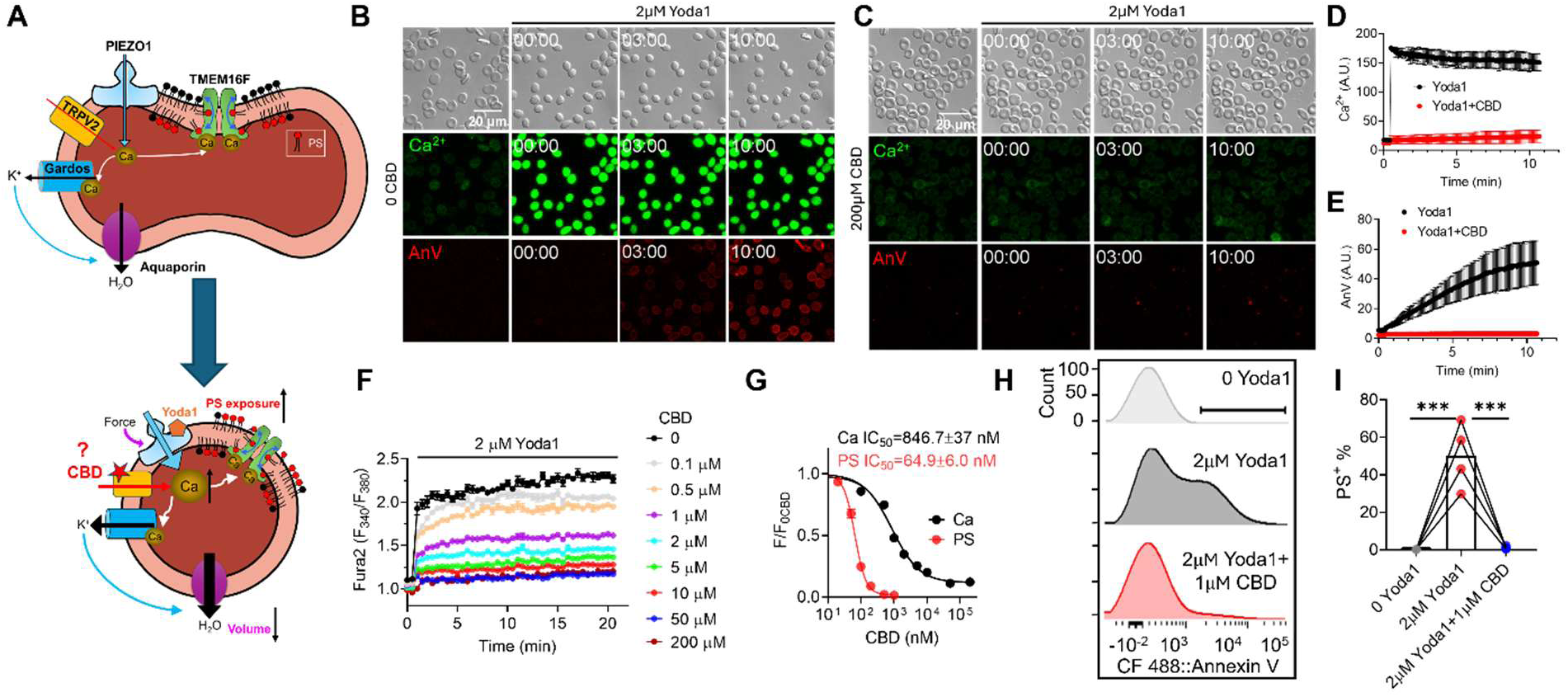
CBD fails to promote Yoda1-induced PS exposure in RBCs. (A) Schematic of the hypothesis: CBD activates TRPV2, promoting Ca²⁺ influx and synergistically enhancing Ca²⁺ entry with PIEZO1. This, in turn, activates TMEM16F and increases PS exposure. (B-C) Yoda1 (2 µM)-induced Ca²⁺ influx (green) and phosphatidylserine (PS) exposure (AnV, red) in human RBCs in the absence (B) and presence (C) of CBD (200 µM). (D-E) Time course of Ca²⁺ influx (D) and lipid scrambling measured by AnV (E) with and without CBD (200 µM). Error bars indicate the standard error of the mean (SEM); N = 3 biological replicates. (F) CBD dose-dependently inhibits Yoda1-induced Ca^2+^ influx in RBCs. Error bars represent ± SEM from 3 replicates. (G) Dose-response curves of CBD inhibition of Ca²⁺ increase in RBCs. Signals were normalized to the 0-CBD condition and fitted using the Hill equation (see *Methods*). (H) Flow cytometry analysis of PS exposure induced by Yoda1 (2 µM) in RBCs with or without CBD (1 µM). RBCs without Yoda1 served as a negative control. (I) Quantification of PS-positive cells from panel H. Each dot represents the average of at least three replicates per blood sample (N = 4 donors). Statistical analysis: unpaired two-sided Student’s *t* test; ***P < 0.001.

To further investigate this unexpected inhibition, we assessed RBC morphology following CBD treatment. In contrast to Yoda1 stimulation, which rapidly dehydrates RBCs into echinocytes through the PIEZO1-Gardos K^+^ channel coupling^12,13^ (Fig. 1A), 200 µM CBD completely prevented Yoda1-induced morphological changes (Fig. S2 and Video S1-2). These observations suggest that CBD, at the concentrations known to activate TRPV2^25–27^, instead inhibits PIEZO1-mediated Ca^2+^ entry in RBCs.

Fura-2 Ca^2+^ imaging revealed that CBD dose-dependently inhibits Yoda1 (2 µM)-induced Ca²⁺ influx in HD RBCs with an IC₅₀ of 0.847 ± 0.037 µM (Fig.1F-G), which is >30-fold lower than the EC_50_ of CBD for human TRPV2^25^. Flow cytometry further demonstrated that CBD potently suppresses Yoda1-induced PS exposure with an IC_50_ of 0.065 ± 0.006 µM, ∼10-fold lower than its IC_50_ for Ca^2+^ entry inhibition (Fig. 1G). Within the range of 0.5-2 µM, CBD completely abolished PS exposure while still allowing PIEZO1-mediated Ca^2+^ entry and RBC morphological changes to occur (Fig. S3). The distinct IC_50_ values for PS exposure and Ca^2+^ entry define a ‘therapeutic window’ in which CBD prevents the detrimental effects of PS exposure while preserving essential PIEZO1 functions in RBCs. Similar ‘therapeutic windows’ were observed with PIEZO1 inhibitors GsMTx4 and benzbromarone in our previous studies^7,13^.

### CBD is a PIEZO1 inhibitor

To determine whether CBD suppresses PS exposure specifically through PIEZO1 inhibition rather than by directly targeting TMEM16F, we examined the effects of 10 µM CBD, a concentration >150-fold higher than the IC_50_ for PS suppression in RBCs, on TMEM16F-mediated lipid scrambling and channel activation in HEK293T cells stably expressed mouse TMEM16F. Lipid scramblase imaging^28^ and patch clamp recording^15,29^ revealed that CBD at this concentration had no detectable effect on ionomycin-induced, TMEM16F-dependent PS exposure (Fig. S4A-C) or Ca^2+^-activated TMEM16F current (Fig. S4D-E). These results indicate that CBD prevents PS exposure in RBCs indirectly by inhibiting upstream PIEZO1 rather than by directly acting on TMEM16F.

To confirm CBD directly inhibits PIEZO1, we used HEK293T cells stably expressing human PIEZO1. Consistent with our observations in HD RBCs (Fig. 1D, F-G), CBD abolished Yoda1-evoked Ca²⁺ influx through heterologously expressed PIEZO1 channels (Fig. 2A-C).

**Figure 2.**
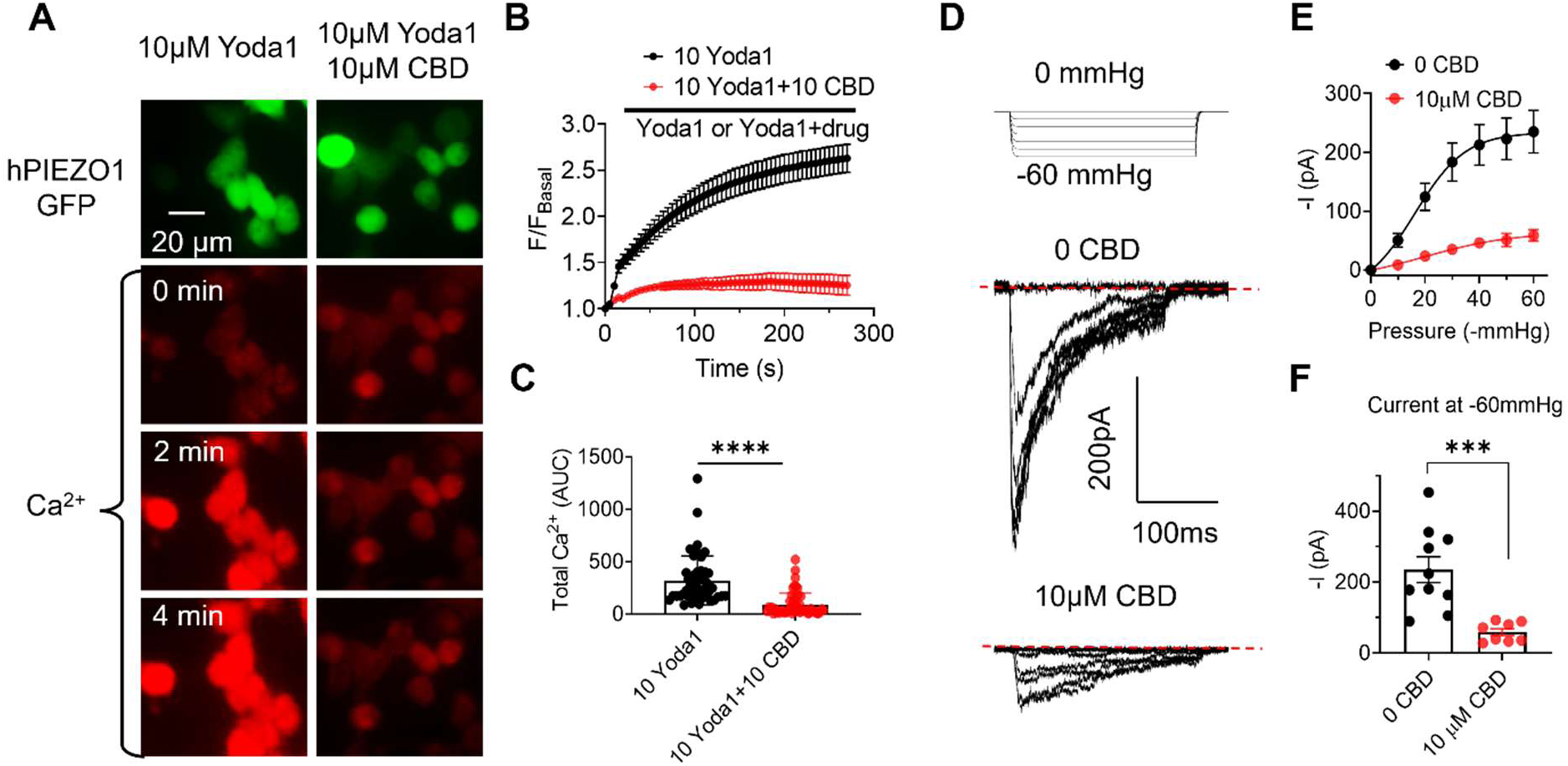
CBD is a PIEZO1 inhibitor. (A) Representative images of Yoda1 (10 µM)-induced Ca²⁺ influx (red) in HEK293 cells overexpressing human PIEZO1 (GFP), in the absence and presence of CBD (10 µM), at the indicated time points. (B) Time course of Yoda1-induced Ca²⁺ influx with or without CBD. Data are presented as mean ± SEM (n = 46 and 52 cells for control and CBD groups, respectively; 3 biological replicates). (C) Quantification of total Ca²⁺ influx, represented as the area under the curve (AUC), with or without CBD (n = 46 and 52 cells for control and CBD groups, respectively; 3 biological replicates). (D) Representative cell-attached patch-clamp recordings of PIEZO1 currents from HEK293T cells in the presence or absence (control) of 10 µM CBD in the pipette solution (extracellular). Currents were elicited by pressure steps from 0 to -60 mmHg at a holding potential of -80 mV. (E) PIEZO1 current-pressure relationships with and without extracellular CBD. Data are shown as mean ± SEM (n = 10 and 8 cells for control and CBD groups, respectively). (F) Comparison of PIEZO1 current amplitudes at -60 mmHg with and without extracellular CBD. Statistical analysis was performed using an unpaired two-sided Student’s t test (***P < 0.001; n = 10 and 8 cells for control and CBD groups, respectively).

Pressure-clamp recordings further demonstrated that 10 µM CBD strongly inhibited PIEZO1 current elicited by mechanical stimulation in cell-attached configuration from the extracellular side (Fig. 2D-F). CBD from the intracellular side also effectively inhibits PIEZO1 current, and the inhibitory effect can be fully washed off (Fig. S5). Notably, it took about 30 seconds to maximize or wash off the CBD effect (Fig. S5B), suggesting that CBD likely intercalates into the lipid bilayer and acts on the transmembrane domain of PIEZO1 to inhibit its opening, similar to its reported modulation of other ion channels^30^.

Taken together, the dose-dependent, reversible inhibition of Yoda1-induced Ca²⁺ influx and mechanically activated PIEZO1 currents by CBD provides strong evidence that CBD directly targets PIEZO1.

### CBD prevents dehydration, hemolysis, PS exposure, and coagulation in HX RBCs

GOF PIEZO1 mutations cause HX, a rare red cell disorder characterized by increased dehydration, hemolysis, iron overload, PS exposure, and thrombotic risk^13,17–19,31–35^. To date, no clinically available PIEZO1 inhibitors have been developed for the treatment of HX. Given the strong translational potential of CBD, we tested whether CBD can inhibit HX-associated mutations and alleviate the related pathological features of human HX RBCs *ex vivo*.

We first examined CBD on PIEZO1-R2488Q, a strong GOF HX mutation^13^, overexpressed in HEK293T cells. CBD not only suppressed Yoda1-evoked Ca²⁺ influx and PS exposure (Fig. S6), but also significantly attenuated mechanically activated PIEZO1 current elicited by pressure clamp stimulation (Fig. 3A-C). These results confirm that CBD effectively inhibits the HX mutant PIEZO1 channels.

**Figure 3.**
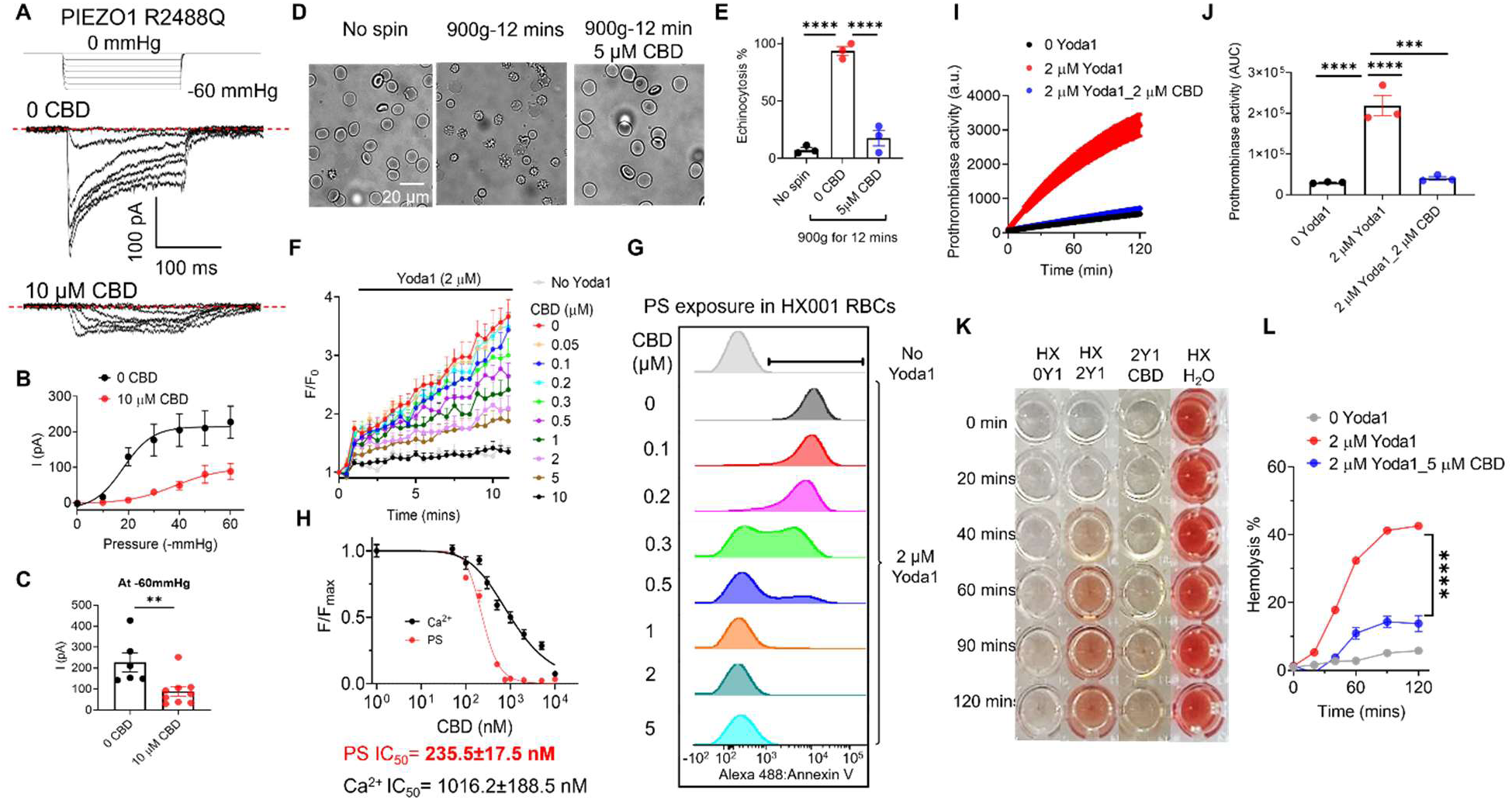
CBD mitigates HX-associated RBC complications by inhibiting PIEZO1. (A) Representative cell-attached patch-clamp recordings of human PIEZO1-R2488Q currents in the presence or absence of 10 µM CBD in the pipette solution. PIEZO1-R2488Q was overexpressed in HEK293T cells. Currents were elicited by pressure steps from 0 to -60 mmHg at a holding potential of −80 mV. (B) PIEZO1-R2488Q current-pressure relationships with and without extracellular CBD. Data are shown as mean ± SEM (n = 6 and 9 cells for the without and with CBD groups, respectively). (C) Comparison of PIEZO1 current amplitudes at -60 mmHg with and without extracellular CBD. Statistical analysis was performed using an unpaired two-sided Student’s t test (**P < 0.01; n = 6 and 9 cells for the without and with CBD groups, respectively). (D) Representative images of HX001 RBCs carrying PIEZO1-R2488Q with and without 5 µM CBD before and after centrifugation. (E) Effect of CBD on centrifugation-induced RBC dehydration (see “Methods” for details). (F) CBD dose-dependently inhibits Yoda1-induced Ca^2+^ influx in HX001 RBCs. Error bars represent ± SEM from 3 replicates. (G) Representative flow cytometry data of CBD inhibition on Yoda1 (2 μM)-induced PS exposure in HX001 RBCs. CF488-AnV was used as a fluorescence PS marker. Each concentration of CBD was repeated in 3 independent experiments. (H) Dose-response curves of CBD inhibition on Ca^2+^ increase (black) and PS exposure (red) in the RBCs from HX001 patient. All signals were normalized to 0 CBD condition and fitted with the Hill equation (see “Methods”). (I) Representative thrombin-mediated fluorogenic results to report prothrombinase activity of HX RBCs under three different conditions: 0 Yoda1, 2 µM Yoda1, and 2 µM Yoda1 with 2 μM CBD. (J) Quantification of prothrombinase activity under the conditions in (I), measured as the area under the curve (AUC). One-way ANOVA followed by Tukey’s test. ***P < 0.001, ****P < 0.0001. Data were obtained from three duplicates for each condition. (K) Representative photos of 2 μM Yoda1(2Y1)-induced hemolysis in HX RBCs with and without 5 μM CBD. 0 Yoda1 (left) and H_2_O (right) serve as negative and positive controls, respectively. (L) Time course of 2 μM Yoda1(Y1)-induced with and without 5 μM CBD in HX001 RBCs. Results were normalized to the water-induced fully lysed group. Yoda1-induced RBCs hemolysis percentage at 120 minutes with and without CBD was compared with an unpaired 2-sided Student *t* test, ∗∗∗∗*P* < .0001, n = 3 repeats for each condition.

Next, we examined whether CBD could prevent force-induced dehydration or echinocytosis in RBCs from an HX patient carrying the PIEZO1-R2488Q mutation. We previously reported that this mutant channel renders HX RBCs’ hypersensitivity to mechanical stimulation, such as centrifugation at 900 g, leading to excessive Ca^2+^ influx, RBC dehydration, and echinocytic morphology of HX RBCs^13^. Treatment with 5 µM CBD, similar to the PIEZO1 inhibitors GsMTx4 and benzbromarone^13^, markedly prevented force-induced echinocytosis (Fig. 3D-E), indicating that CBD protects HX RBCs from dehydration and preserves their mechanical stability.

Enhanced PIEZO1-TMEM16F coupling and the resulting PS exposure are additional hallmarks of HX RBCs, contributing to anemia, splenomegaly, and post-splenectomy thrombosis^13^. We therefore assessed whether CBD could reduce this abnormal PS exposure. CBD dose-dependently suppressed Yoda1-induced Ca²⁺ influx (IC₅₀ = 1.016 ± 0.189 µM; Fig. 3F, 3H) and PS exposure (IC₅₀ = 0.236 ± 0.018 µM; Fig. 3G-H). Notably, CBD maintained a clear ‘therapeutic window’ in HX RBCs, where sub-µM concentrations potently reduce PS exposure while preserving partial PIEZO1 activity (Fig. 3H). Although the IC_50_ values for HX RBCs are higher than those of HD RBCs due to PIEZO1 GOF, sub-micromolar CBD remains sufficient to suppress PS exposure in the HX RBCs.

We recently postulated that enhanced PIEZO1-TMEM16F coupling induced PS exposure in HX RBCs is likely a major contributor to heightened thrombotic risk, particularly after splenectomy^36^, when PS⁺ RBCs are no longer efficiently cleared. Consistent with this, 2 µM Yoda1 triggered robust thrombin generation in HX RBCs as measured by a fluorescent prothrombin assay (Figs. 3I-J). Importantly, 2 µM CBD completely abolished Yoda1-induced thrombin generation, suggesting that PIEZO1 inhibition by CBD could mitigate the thrombotic risk associated with HX.

Finally, we examined whether CBD could prevent PIEZO1 hyperactivation-induced hemolysis, a likely contributor to intravascular hemolysis and iron overload commonly observed in PIEZO1-dependent HX patients^36,37^. Consistent with prior findings^13^, 2 µM Yoda1 induced time-dependent hemolysis of the HX RBCs *ex vivo*, while 5 µM CBD almost completely prevented it (Fig. 3K-L).

Collectively, these results demonstrate that CBD, as a potent inhibitor of PIEZO1, effectively suppresses abnormal Ca²⁺ influx, PS exposure, hypercoagulation, dehydration, and hemolysis in HX RBCs, underscoring its potent inhibitory action on PIEZO1 GOF activity. The sub-micromolar potency of CBD in preventing PS exposure and downstream complications highlights its strong translational potential as a candidate therapeutic or lead compound for HX and related PIEZO1-mediated disorders.

### CBD restores Ca^2+^ and membrane homeostasis in sickle (SS) RBCs

Aberrant activation of PIEZO1 contributes to both basal and deoxygenation-induced Ca²⁺ elevation in SS RBCs and enhances PIEZO1-TMEM16F coupling, promoting excessive PS exposure, PS+ microparticle release, hypercoagulability, and RBC-endothelium adhesion^7^. We therefore hypothesized that by inhibiting PIEZO1, CBD could attenuate these pathological processes and restore calcium and membrane homeostasis in SCD RBCs. Indeed, 2 µM CBD robustly suppressed deoxygenation-induced Ca²⁺ elevation in SS RBCs (Fig. 4A). Consequently, deoxygenation-induced PS exposure and PS^+^ microparticle release under deoxygenation were both reduced by CBD to near-normoxic levels (Fig. 4B-C). When deoxygenated, irreversibly sickled RBCs were challenged with Yoda1 to mimic mechanical stimulations in circulation^7^, we observed a rapid PIEZO1-mediated Ca²⁺ influx that was inhibited by CBD in a concentration-dependent manner (Fig. 4D, 4F). Flow cytometry confirmed that CBD dose-dependently reduced Yoda1-induced PS exposure in sickled RBCs (Figs. 4E-F).

**Figure 4.**
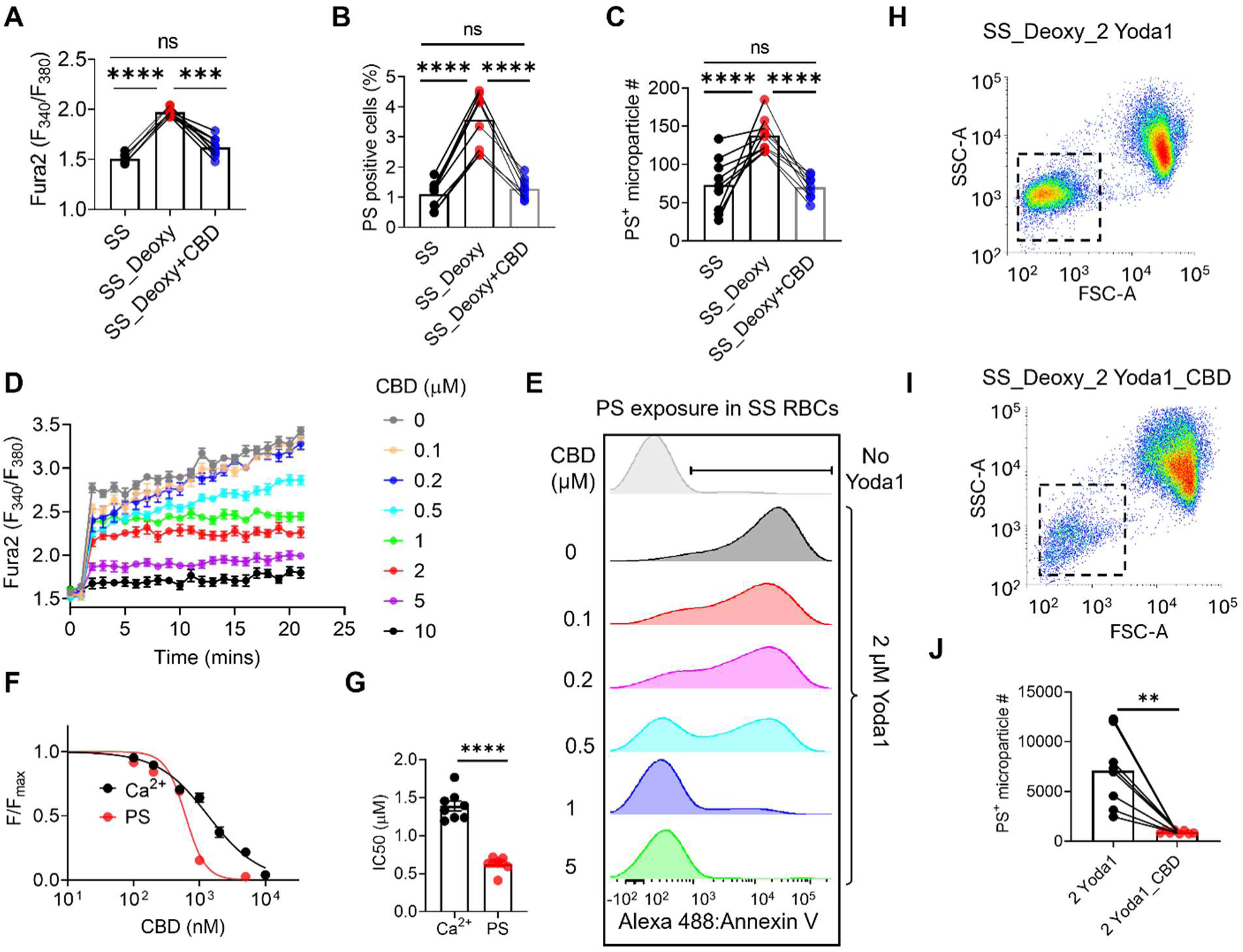
CBD reduces basal and Yoda1-induced Ca²⁺ levels, PS⁺ cells, and PS⁺ microparticles in sickled RBCs. (A) Basal intracellular Ca²⁺ levels in red blood cells (RBCs) from sickle cell disease (SS) patients under normoxic and deoxygenated (Deoxy) conditions with or without 2 μM CBD. One-way ANOVA with Tukey’s test. ***P < 0.001, ****P < 0.0001, ns = not significant. n = 9 patients; data represent triplicate means per sample. (B) Percentage of PS⁺ cells under the same conditions as in (A). One-way ANOVA with Tukey’s test. ****P < 0.0001, ns = not significant. n = 9 patients; data represent triplicate means per sample. (C) Quantification of PS⁺ microparticles under the same conditions as in (A). One-way ANOVA with Tukey’s test. ****P < 0.0001, ns = not significant. n = 9 patients; data represent triplicate means per sample. (D) Fura-2 measurements of Yoda1-induced Ca²⁺ influx in deoxygenated SS RBCs at different CBD concentrations. n = 3 patients; data are mean ± SEM from triplicates. (E) Representative flow cytometry analysis of CBD inhibition of Yoda1-induced PS exposure in deoxygenated SS RBCs. Each CBD concentration was tested in 3 independent experiments. (F) Dose-response curves of CBD inhibition of Yoda1-induced Ca²⁺ influx (black) and PS exposure (red) in deoxygenated SS RBCs. Data were normalized to the 0 μM CBD condition and fitted with the Hill equation. (G) IC₅₀ values for CBD inhibition of Yoda1-induced Ca²⁺ influx and PS exposure in deoxygenated SS RBCs. Unpaired two-sided Student’s t test. **P < 0.01. n = 8 patients; data represent triplicate means per sample. (H-I) Representative flow cytometry plots of microparticle release (dotted rectangles) induced by 2 μM Yoda1 in deoxygenated SS RBCs without (H) or with (I) 1 μM CBD. (J) Comparison of PS⁺ microparticle release from deoxygenated SS RBCs treated with 2 μM Yoda1 in the absence or presence of 1 μM CBD. Paired two-sided Student’s t-test. **P < 0.01. n = 8 patients; data represent triplicate means per sample.

As in HX RBCs (Figs. 3F-H), CBD displayed higher apparent potency in preventing PS exposure than in blocking PIEZO1-mediated Ca²⁺ entry (Fig. 4G), defining a ‘therapeutic window’ where detrimental PS externalization is suppressed while essential PIEZO1 functions are preserved (Fig. 4F). In addition, 1 µM CBD significantly reduced Yoda1-induced PS^+^ microparticle generation from deoxygenated SS RBCs (Fig. 4H-J). These findings demonstrate that CBD potently inhibits aberrant PIEZO1 activation and subsequent TMEM16F activation in sickled RBCs, thereby restoring Ca^2+^ and membrane homeostasis.

### CBD prevents PIEZO1-driven sickling, coagulation, and endothelial adhesion in SS RBCs

We next evaluated the effects of CBD on mitigating PIEZO1-driven complications in SCD. Treatment with 10 μM CBD effectively prevented deoxygenation-induced sickling of SS RBCs (Fig. 5A-B), indicating a clear anti-sickling effect. This protection likely arises from CBD’s inhibition of PIEZO1, as enhanced PIEZO1 activity in SS RBCs exacerbates sickling through PIEZO1-Gardos K⁺ channel coupling, which promotes RBC dehydration and markedly increases the propensity for sickling^8^.

**Figure 5.**
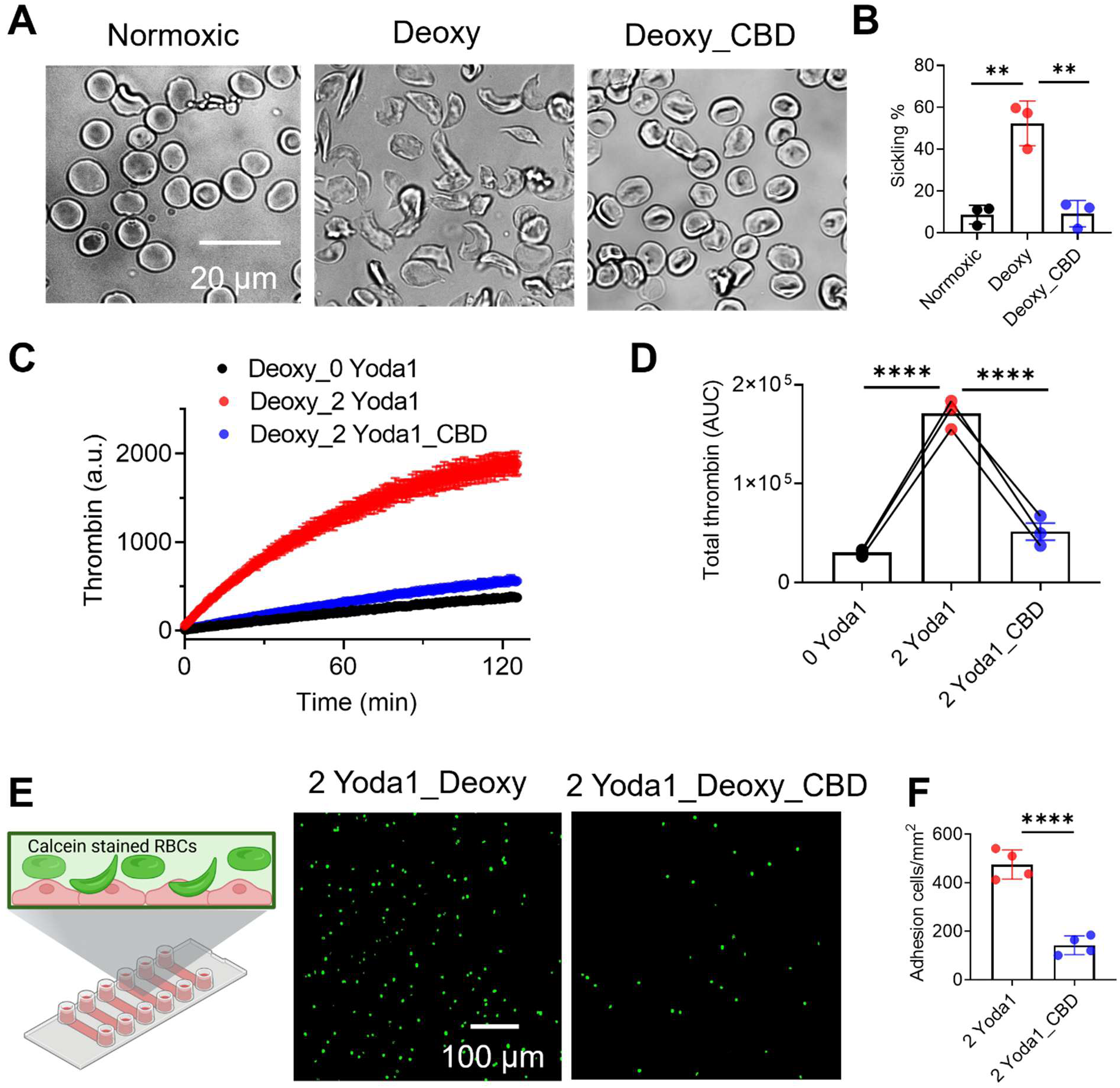
CBD mitigates SCD-associated RBC complications in vitro. (A) Representative images of SS RBC morphology under normoxic and deoxygenated conditions without or with 10 μM CBD. (B) Quantification of sickle-shaped cells under the conditions in (A). One-way ANOVA with Tukey’s test. **P < 0.01. n = 3 patients; data represent triplicate means per sample. (C) Representative thrombin-mediated fluorogenic readouts reporting prothrombinase activity of SS RBCs under three conditions. (D) Quantification of prothrombinase activity in (C), measured as area under the curve (AUC). One-way ANOVA with Tukey’s test. ****P < 0.0001. Data from 3 SCD patients, with duplicates for each sample. (E) Representative fluorescence images of SS RBC adhesion to HUVEC endothelial cells (Left: schematic). Calcein-loaded SS RBCs (green) were deoxygenated with 2 μM Yoda1 or 2 μM Yoda1 +2 μM CBD. (F) Quantification of adhered SS RBCs to HUVEC under the conditions in (E). Unpaired two-sided Student’s t test. ****P < 0.0001. n = 4 patients; data represent triplicate means per sample.

Hypercoagulability, a hallmark of SCD, elevates the risks of thrombosis and vaso-occlusion^38,39^. Because PS⁺ RBCs and microparticles play key roles in promoting coagulation, we next tested whether CBD could suppress thrombin generation in SS RBCs using a fluorogenic thrombin generation assay^7^. Under deoxygenated conditions, Yoda1 markedly increased thrombin generation compared with deoxygenation alone, whereas co-treatment with 2 μM CBD substantially blunted this response (Fig. 5C-D). These results are consistent with CBD’s ability to disrupt PIEZO1-TMEM16F coupling and thereby reduce PS exposure.

Increased PIEZO1 activity and its coupling to TMEM16F also enhance SS RBC adhesion to endothelial cells^7,8,40^. To assess whether CBD can alleviate this effect, we quantified RBC-endothelium interactions using calcein-labeled SS RBCs^7^. Deoxygenated and Yoda1-stimulated SS RBCs exhibited robust endothelial adhesion, which was markedly reduced by 2 μM CBD (Fig. 5E-F).

In summary, CBD significantly reduces sickling, thrombin generation, and endothelial adhesion, three key pathogenic processes in SCD. These findings highlight that CBD’s therapeutic potential extends beyond its established role in pain management, positioning it as a promising mechanistically targeted agent for mitigating PIEZO1-driven SCD complications.

### CBD inhibits PIEZO2-mediated touch sensation

CBD has been widely used to mitigate hyperalgesia associated with various diseases, including SCD^24^. PIEZO2, a close structural homolog of PIEZO1, is abundantly expressed in sensory neurons, where it mediates light touch, proprioception, and mechanical pain^41,42^. Given CBD’s inhibitory effect on PIEZO1, we hypothesized that CBD might also inhibit PIEZO2 activity. To test this, we measured shear stress-evoked Ca²⁺ responses in PIEZO1-knockout HEK293T cells overexpressing human PIEZO2. Under control conditions, shear stress (27.2 dyn/cm²) elicited robust Ca²⁺ influx, as measured by Ca^2+^ imaging (Fig. 6A). Co-application of 10 µM CBD markedly reduced both the amplitude and total Ca²⁺ influx (Figs. 6B-C). Whole-cell patch-clamp recordings further showed that shear stress-evoked inward currents were strongly suppressed by 10 µM CBD (Figs. 6D-E), demonstrating direct inhibition of PIEZO2 channel activity by CBD.

**Figure 6.**
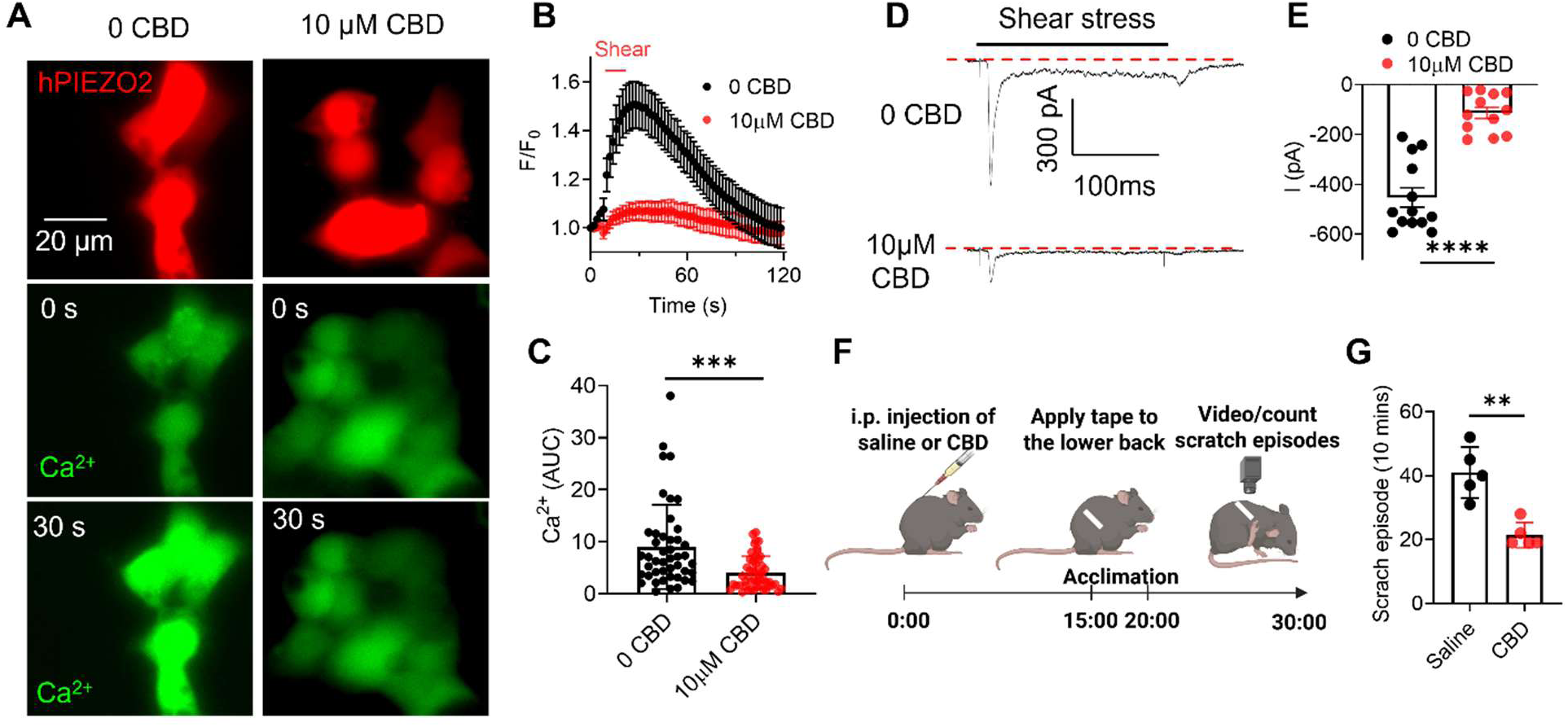
CBD blocks PIEZO2. (A) Representative images of shear stress (27.2 dyn/cm²)-induced Ca²⁺ influx (green) in HEK293T cells overexpressing human PIEZO2 (mCherry), in the absence (left) or presence (right) of 10 μM CBD, at the indicated time points. (B) Time course of shear stress-induced Ca²⁺ influx with or without CBD. Data are mean ± SEM (n = 45 and 50 cells for 0 and 10 μM CBD groups, respectively; 3 biological replicates). (C) Quantification of total Ca²⁺ influx, expressed as area under the curve (AUC), with or without CBD. Data are mean ± SEM (n = 45 and 50 cells for 0 and 10 μM CBD groups, respectively; 3 biological replicates). (D) Representative whole-cell patch-clamp recordings of human PIEZO2 currents with 0 or 10 μM CBD in the pipette solution. Human PIEZO2 was overexpressed in PIEZO1 KO HEK293T cells. Currents were elicited by shear stress (27.2 dyn/cm²) at a holding potential of -80 mV. (E) Quantification of PIEZO2 currents with and without extracellular CBD. Data are mean ± SEM. Unpaired two-sided Student’s t test, ****P < 0.0001 (n = 13 and 12 cells for 0 and 10 μM CBD groups, respectively). (F) Schematic of the tape response assay (see *Methods* for details). (G) Quantification of scratch episodes in saline control versus CBD-treated mice (20 mg/kg, i.p.). Unpaired two-sided Student’s t test, **P < 0.01 (n = 5 mice per group).

To evaluate physiological relevance *in vivo*, we employed the tape response assay (Fig. 6F), which assesses tactile sensation in mice^43^. In this test, control mice sensed the tape attached to their back and attempted to remove it by repetitive scratching, reflecting intact PIEZO2-mediated mechanosensation (Fig. 6G, Video S3). In contrast, CBD-treated mice (20 mg/kg, i.p.) exhibited significantly fewer scratching episodes than saline-treated controls, consistent with dampened mechanosensory signaling (Fig. 6G, Video S4). Collectively, these findings demonstrate that CBD inhibits PIEZO2 activity both *in vitro* and *in vivo*, providing a plausible mechanism for its reduction of mechanical hypersensitivity and pain associated with SCD and other disorders involving PIEZO2-mediated sensory pathways.

## DISCUSSION

This study reveals CBD as an unexpected, mechanistically targeted inhibitor of PIEZO1 channels that restores Ca^2+^ and membrane homeostasis in diseased RBCs. This reframes CBD from a symptomatic analgesic to a potential disease-modifying agent for red cell disorders. CBD directly inhibits PIEZO1 without detectable effect on TMEM16F scramblase, thereby disengaging the PIEZO-TMEM16F axis that drives PS exposure and its downstream procoagulant and adhesive complications. Functionally, CBD prevents force-induced dehydration, limits hemolysis, and suppresses thrombin generation in HX RBCs bearing PIEZO1 GOF mutations; and in SCD RBCs, it reduces deoxygenation- and Yoda1-evoked Ca²⁺ influx, lowers PS exposure and PS⁺ microparticle shedding, decreases sickling, and diminishes endothelial adhesion. Given CBD’s wide availability and favorable safety profile^44^, identifying CBD as a potent PIEZO1 inhibitor in RBCs provides a new mechanistic foundation for clinical evaluation in SCD. This approach, by targeting core RBC pathobiology rather than pain alone, will motivate the development of PIEZO-selective CBD analogs with optimized efficacy and safety.

Excessive PS exposure is a key pathological driver of red cell disorders, including SCD and HX^7,13^. A notable feature of our dose-response analysis is a therapeutic window in which partial inhibition of PIEZO1 disengages downstream TMEM16F without abolishing essential PIEZO1 functions. Sub-micromolar CBD, which is readily achieved clinically in epilepsy patients at approved doses^21,45,46^, potently suppresses PS exposure while only partially reducing PIEZO1-mediated Ca²⁺ entry and preserving RBC shape/volume responses to mechanical or chemical stimuli to PIEZO1. This window is mechanistically coherent. TMEM16F scramblase requires higher/spatially sustained Ca²⁺ signals^15^ than those needed for some baseline PIEZO1-dependent functions, such as Gardos channel-mediated volume adjustments^12,47^. Therefore, modest attenuation of Ca²⁺ influx can selectively inhibit the scramblase without abolishing entire PIEZO1 activity. Given that PS exposure is a key pathophysiological driver in HX^13^ and SCD^7,9,40,48^, this window also has high clinical relevance. We previously observed analogous windows with GsMTx4 and benzbromarone^7,13^. But, CBD uniquely combines this pharmacological profile with broad human exposure data and a favorable safety record at clinically used doses^21–23,44,45,49^, strengthening its translational potential as a disease-modifying therapy for HX and SCD.

Clinical evidence and recent SCD mouse studies support CBD’s beneficial effects in mitigating SCD complications, such as reducing vaso-occlusive crisis-related hospitalizations in patients and decreasing hyperalgesia and inflammation in mice^24,50,51^. Given growing evidence that PIEZO1 functions as a critical mechanosensor in immune cells, including regulating activation, migration, and cytokine production^52–54^, it is plausible that CBD also inhibits PIEZO1 in these cells and attenuates inflammation in SCD. Beyond erythrocytes, we found that CBD blocks PIEZO2 *in vitro* and dampens touch-evoked responses *in vivo*. Because PIEZO2 is the principal mechanotransduction channel for light touch and contributes to mechanical pain, these findings provide a previously unrecognized molecular mechanism for CBD’s effects on mechanical hypersensitivity, including in SCD. With the identification of CBD as a potent PIEZO channel inhibitor, future clinical and preclinical studies should define dose ranges, exposure-response relationships, and long-term benefit-risk profiles that maximize RBC benefits.

Despite that PIEZO1 GOF is well-known as the causal lesion of HX^17^, no PIEZO1-directed therapy currently exists. CBD thus offers a direct etiologic, translational strategy to mitigate dehydration, hemolysis, iron overload, and hypercoagulability, including the increased post-splenectomy thrombotic risk. Future studies in mouse models and early-phase trials are needed to determine whether clinically relevant CBD exposures can chronically normalize aberrant PIEZO1 activity in RBCs and reduce hemolysis, iron overload, and thrombin generation. The identification of CBD as a PIEZO1 inhibitor thus provides a possible solution for the urgent unmet need in treating this rare disorder.

Beyond red cell disorders, enhanced PIEZO1/PIEZO2 activity is implicated in diverse pathologies, including vascular disease (arteriovenous malformations^55^, atherosclerosis^56^), cardiac remodeling^57–59^ (hypertrophy, fibrosis), osteoarthritis^60,61^, developmental skeletal disorders^62,63^ (distal arthrogryposis, Marden–Walker syndrome), cancer mechanotransduction^64,65^, and sensory/proprioceptive abnormalities^41,66–69^. Therefore, identifying CBD as a small-molecule PIEZO inhibitor will also motivate targeted preclinical and clinical studies across these systems and supports the development of PIEZO-selective CBD analogs to maximize efficacy while minimizing sensory side effects.

Mechanistically, CBD’s bidirectional membrane access and the slow onset/offset observed in patch-clamp experiments suggest that CBD intercalates into the plasma membrane and inhibits PIEZO channels by interacting with transmembrane/lipid-facing regions. Structure-function studies, including mutational scanning, cryo-EM with bound ligands, and molecular docking and simulation, will be essential to pinpoint CBD interaction sites and guide the design of more selective and potent PIEZO inhibitors for red cell disorders.

In summary, CBD emerges as a PIEZO1 inhibitor that interrupts the PIEZO1-TMEM16F coupling, reduces PS-driven coagulant and adhesive phenotypes, and lessens sickling, key pathogenic processes in HX and SCD, while also dampening PIEZO2-mediated mechanosensation. These findings provide a strong rationale to clinically evaluate CBD and to develop PIEZO-selective CBD analogs as disease-modifying therapies for red cell disorders and beyond.

## METHODS

### Human subjects

All human studies were approved by the Institutional Review Board at Duke University (IRB# Pro00007816, Pro00109511, and Pro00104933). Written informed consent was obtained from healthy donors, HX and SCD patients. Our previous study shows that HX001 carries the PIEZO1 p. Arg2488Gln (c.7463G>A) mutation in a heterozygous state^13^.

### Human red blood cell collection

Human blood was collected into tubes containing acid-citrate-dextrose (ACD) buffer and centrifuged at 200 × *g* for 12 min. To minimize mechanical activation, hereditary xerocytosis (HX) red blood cells (RBCs) were centrifuged at 100 × *g* for 12 min. After removal of the supernatant, RBC pellets were washed 2-3 times with phosphate-buffered saline (PBS) using the same centrifugation-resuspension protocol.

### Deoxygenation to induce sickling

Sodium metabisulfite was used to induce red blood cell (RBC) sickling as previously described ^7^. To limit oxidative damage^70^, a fresh 0.2% sodium metabisulfite solution was prepared immediately before use. Packed RBCs (100 µL) were added to 1.5 mL of the solution in 1.5-mL microcentrifuge tubes, which were then sealed with pre-warmed sealing wax to prevent gas exchange. Samples were incubated at 37 °C for 2 h to promote irreversible sickling.

### Mice

Mouse handling and usage were carried out in strict compliance with the protocol approved by the Institutional Animal Care and Use Committee at Duke University, in accordance with National Institute of Health guidelines. Adult C57/BL6J mice of both sexes were used for the behavioral test.

### Tape response assay

A modified tape-on-paw assay was performed ^43^. Mice were acclimated in circular Plexiglas chambers for 5 min. A 3-cm strip of standard laboratory tape was gently applied to the dorsal skin of the mice to ensure firm adhesion. Behavior was recorded for 10 min, and scratch episodes directed toward the tape were counted. A “response” was scored for either (i) a pause in normal activity followed by biting/scratching directed at the tape or (ii) a characteristic “wet-dog shake” aimed at dislodging the tape.

### Statistical analysis

All statistical analyses were performed with Clampfit 10 (Molecular Devices), Excel or Prism software (GraphPad). Two-tailed Student’s t-test was used for single comparisons between two groups (paired or unpaired), and one-way ANOVA followed by Tukey’s test was used for multiple comparisons. Data were represented as mean ± standard error of the mean (SEM) unless stated otherwise.

Other methods are described in the supplemental Methods.

## Data availability

All study data are included in the article and/or SI Appendix. Contact Dr. Huanghe Yang for data and material requests at.

## ACKNOWLEDGEMENT

We thank Alejandra Gonzalez Torres and Ryan Chou for their technical assistance in this study. This work was supported by Duke University (to HY), American Society of Hematology (ASH) Scholar Award (to PL, 2024–2026) and ASH Graduate Hematology Award (to YSW, 2023–2025).

## AUTHORSHIP

Contributions: Conceptualization: HY; Methodology: PL, YSW, KZS, YZ, MD, MD, SK, HY; Investigation: PL, YSW, KZS; Data curation and analysis: PL, YSW, KZS; Visualization: PL, YSW, KZS, HY; Validation: PL, YSW, KZS; Writing – original draft: HY, PL; Writing – review & editing: all authors; Resources: HY, GMA, MJT; Supervision and project administration: HY; Funding acquisition: HY, PL, YSW.

## References

1. Bunn HF. Pathogenesis and treatment of sickle cell disease. N Engl J Med. 1997;337(11):762–769.

2. Manodori AB, Barabino GA, Lubin BH, Kuypers FA. Adherence of phosphatidylserine-exposing erythrocytes to endothelial matrix thrombospondin. Blood. 2000;95(4):1293–1300.

3. Telen MJ, Malik P, Vercellotti GM. Therapeutic strategies for sickle cell disease: towards a multi-agent approach. Nat Rev Drug Discov. 2019;18(2):139–158.

4. Barak M, Hu C, Matthews A, Fortenberry YM. Current and Future Therapeutics for Treating Patients with Sickle Cell Disease. Cells. 2024;13(10).

5. Cavazzana M, Corsia A, Brusson M, Miccio A, Semeraro M. Treating Sickle Cell Disease: Gene Therapy Approaches. Annu Rev Pharmacol Toxicol. 2024.

6. DeMartino PC, Haag MB, Caughey AB, Roth JA. A budget impact analysis of gene therapy for sickle cell disease: an updated analysis. Blood Adv. 2024;8(17):4658–4661.

7. Liang P, Wan Y-CS, Shan KZ, et al. Targeting PIEZO1-TMEM16F Coupling to Mitigate Sickle Cell Disease Complications. American Journal of Hematology. 2025;n/a(n/a).

8. Nader E, Conran N, Leonardo FC, et al. Piezo1 activation augments sickling propensity and the adhesive properties of sickle red blood cells in a calcium-dependent manner. Br J Haematol. 2023.

9. Wadud R, Hannemann A, Rees DC, Brewin JN, Gibson JS. Yoda1 and phosphatidylserine exposure in red cells from patients with sickle cell anaemia. Scientific Reports. 2020;10(1).

10. Romero LO, Bade M, Elsherif L, et al. Enhanced PIEZO1 function contributes to the pathogenesis of sickle cell disease. Proc Natl Acad Sci U S A. 2025;122(40):e2514863122.

11. Rooks H, Brewin J, Gardner K, et al. A gain of function variant in PIEZO1 (E756del) and sickle cell disease. Haematologica. 2019;104(3):e91–e93.

12. Cahalan SM, Lukacs V, Ranade SS, Chien S, Bandell M, Patapoutian A. Piezo1 links mechanical forces to red blood cell volume. eLife. 2015;4.

13. Liang P, Zhang Y, Wan YCS, et al. Deciphering and disrupting PIEZO1-TMEM16F interplay in hereditary xerocytosis. Blood. 2024;143(4):357–369.

14. Vaisey G, Banerjee P, North AJ, Haselwandter CA, MacKinnon R. Piezo1 as a force-through-membrane sensor in red blood cells. Elife. 2022;11.

15. Yang H, Kim A, David T, et al. TMEM16F forms a Ca2+-activated cation channel required for lipid scrambling in platelets during blood coagulation. Cell. 2012;151(1):111–122.

16. Suzuki J, Umeda M, Sims PJ, Nagata S. Calcium-dependent phospholipid scrambling by TMEM16F. Nature. 2010;468(7325):834–838.

17. Gallagher PG. Disorders of erythrocyte hydration. Blood. 2017;130(25):2699–2708.

18. Ma S, Cahalan S, Lamonte G, Winzeler EA, Andersen KG, Patapoutian A. Common PIEZO1 Allele in African Populations Causes RBC Dehydration and Attenuates Plasmodium Infection. Cell. 2018;173:443–455.e412.

19. Glogowska E, Schneider ER, Maksimova Y, et al. Novel mechanisms of PIEZO1 dysfunction in hereditary xerocytosis. Blood. 2017;130(16):1845–1856.

20. Azevedo VF, Kos IA, Vargas-Santos AB, da Rocha Castelar Pinheiro G, Dos Santos Paiva E. Benzbromarone in the treatment of gout. Adv Rheumatol. 2019;59(1):37.

21. Moreira FA, de Oliveira ACP, Santos VR, Moraes MFD. Cannabidiol and epilepsy. Int Rev Neurobiol. 2024;177:135–147.

22. Mohammed SYM, Leis K, Mercado RE, Castillo MMS, Miranda KJ, Carandang RR. Effectiveness of Cannabidiol to Manage Chronic Pain: A Systematic Review. Pain Manag Nurs. 2024;25(2):e76–e86.

23. Nascimento GC, Escobar-Espinal D, Balico GG, Silva NR, Del-Bel E. Cannabidiol and pain. Int Rev Neurobiol. 2024;177:29–63.

24. Cherukury HM, Argueta DA, Garcia N, et al. Cannabidiol attenuates hyperalgesia in a mouse model of sickle cell disease. Blood. 2023;141(2):203–208.

25. Pumroy RA, Samanta A, Liu Y, et al. Molecular mechanism of TRPV2 channel modulation by cannabidiol. eLife. 2019;8:e48792.

26. Belkacemi A, Trost CF, Tinschert R, et al. The TRPV2 channel mediates Ca2+ influx and the Δ9-THC-dependent decrease in osmotic fragility in red blood cells. Haematologica. 2021;106(8):2246–2250.

27. Egée S, Kaestner L. The Transient Receptor Potential Vanilloid Type 2 (TRPV2) Channel-A New Druggable Ca(2+) Pathway in Red Cells, Implications for Red Cell Ion Homeostasis. Front Physiol. 2021;12:677573.

28. Le T, Jia Z, Le SC, Zhang Y, Chen J, Yang H. An inner activation gate controls TMEM16F phospholipid scrambling. Nature Communications. 2019;10:1846.

29. Liang P, Yang H. Molecular underpinning of intracellular pH regulation on TMEM16F. J Gen Physiol. 2021;153(2).

30. Watkins AR. Cannabinoid interactions with ion channels and receptors. Channels (Austin). 2019;13(1):162–167.

31. Jankovsky N, Caulier A, Demagny J, et al. Recent advances in the pathophysiology of PIEZO1-related hereditary xerocytosis. Am J Hematol. 2021;96(8):1017–1026.

32. Zarychanski R, Schulz VP, Houston BL, et al. Mutations in the mechanotransduction protein PIEZO1 are associated with hereditary xerocytosis. Blood. 2012;120(9):1908–1915.

33. Albuisson J, Murthy SE, Bandell M, et al. Dehydrated hereditary stomatocytosis linked to gain-of-function mutations in mechanically activated PIEZO1 ion channels. Nature Communications. 2013;4:1884.

34. Andolfo I, Alper SL, De Franceschi L, et al. Multiple clinical forms of dehydrated hereditary stomatocytosis arise from mutations in PIEZO1. Blood. 2013;121:3925–3935.

35. Ma S, Dubin AE, Zhang Y, et al. A role of PIEZO1 in iron metabolism in mice and humans. Cell. 2021;184(4):969–982 e913.

36. Picard V, Guitton C, Thuret I, et al. Clinical and biological features in PIEZO1-hereditary xerocytosis and gardos channelopathy: A retrospective series of 126 patients. Haematologica. 2019;104:1554–1564.

37. Archer NM, Shmukler BE, Andolfo I, et al. Hereditary xerocytosis revisited. Am J Hematol. 2014;89(12):1142–1146.

38. van Tits LJ, van Heerde WL, Landburg PP, et al. Plasma annexin A5 and microparticle phosphatidylserine levels are elevated in sickle cell disease and increase further during painful crisis. Biochemical and Biophysical Research Communications. 2009;390:161–164.

39. Whelihan MF, Lim MY, Mooberry MJ, et al. Thrombin generation and cell-dependent hypercoagulability in sickle cell disease. J Thromb Haemost. 2016;14(10):1941–1952.

40. Setty BN, Kulkarni S, Stuart MJ. Role of erythrocyte phosphatidylserine in sickle red cell-endothelial adhesion. Blood. 2002;99(5):1564–1571.

41. Murthy SE, Loud MC, Daou I, et al. The mechanosensitive ion channel Piezo2 mediates sensitivity to mechanical pain in mice. Science Translational Medicine. 2018;10(462):eaat9897.

42. Dubin Adrienne E, Schmidt M, Mathur J, et al. Inflammatory Signals Enhance Piezo2-Mediated Mechanosensitive Currents. Cell Reports. 2012;2(3):511–517.

43. Ranade SS, Woo S-H, Dubin AE, et al. Piezo2 is the major transducer of mechanical forces for touch sensation in mice. Nature. 2014;516:121–125.

44. Kicman A, Toczek M. The Effects of Cannabidiol, a Non-Intoxicating Compound of Cannabis, on the Cardiovascular System in Health and Disease. Int J Mol Sci. 2020;21(18).

45. Cafaro A, Riva A, Pigliasco F, et al. Long-term plasma monitoring of THC and CBD in pediatric drug-resistant epilepsy: Implications for cannabidiol therapy with Epidyolex(R). Epilepsia Open. 2025;10(5):1699–1704.

46. Contin M, Mohamed S, Santucci M, et al. Cannabidiol in Pharmacoresistant Epilepsy: Clinical Pharmacokinetic Data From an Expanded Access Program. Frontiers in Pharmacology. 2021;Volume 12 - 2021.

47. Bogdanova A, Makhro A, Wang J, Lipp P, Kaestner L. Calcium in red blood cells-a perilous balance. Int J Mol Sci. 2013;14(5):9848–9872.

48. Wood BL, Gibson DF, Tait JF. Increased erythrocyte phosphatidylserine exposure in sickle cell disease: flow-cytometric measurement and clinical associations. Blood. 1996;88(5):1873–1880.

49. Millar SA, Stone NL, Bellman ZD, Yates AS, England TJ, O’Sullivan SE. A systematic review of cannabidiol dosing in clinical populations. Br J Clin Pharmacol. 2019;85(9):1888–1900.

50. Mayrand L, Tarbé de Saint Hardouin A-L, Maciel TT, et al. Dramatic efficacy of cannabidiol on refractory chronic pain in an adolescent with sickle cell disease. American Journal of Hematology. 2023;98(11):E295–E297.

51. Curtis SA, Brandow AM, DeVeaux M, Zeltermam D, Devine L, Roberts JD. Daily Cannabis Users with Sickle Cell Disease Show Fewer Admissions than Others with Similar Pain Complaints. Cannabis and Cannabinoid Research. 2020;5(3):255–262.

52. Aykut B, Chen R, Kim JI, et al. Targeting Piezo1 unleashes innate immunity against cancer and infectious disease. Sci Immunol. 2020;5(50).

53. Solis AG, Bielecki P, Steach HR, et al. Mechanosensation of cyclical force by PIEZO1 is essential for innate immunity. Nature. 2019;573(7772):69–74.

54. Qu P, Zhang H. The dual role of Piezo1 in tumor cells and immune cells: a new target for cancer therapy. Front Immunol. 2025;16:1635388.

55. Park H, Lee S, Furtado J, et al. PIEZO1 Overexpression in Hereditary Hemorrhagic Telangiectasia Arteriovenous Malformations. Circulation. 2025;152(9):599–615.

56. Lan Y, Lu J, Zhang S, et al. Piezo1-Mediated Mechanotransduction Contributes to Disturbed Flow-Induced Atherosclerotic Endothelial Inflammation. J Am Heart Assoc. 2024;13(21):e035558.

57. Bartoli F, Evans EL, Blythe NM, et al. Global PIEZO1 Gain-of-Function Mutation Causes Cardiac Hypertrophy and Fibrosis in Mice. Cells. 2022;11(7).

58. Zhang Y, Su SA, Li W, et al. Piezo1-Mediated Mechanotransduction Promotes Cardiac Hypertrophy by Impairing Calcium Homeostasis to Activate Calpain/Calcineurin Signaling. Hypertension. 2021;78(3):647–660.

59. Jiang F, Yin K, Wu K, et al. The mechanosensitive Piezo1 channel mediates heart mechano-chemo transduction. Nat Commun. 2021;12(1):869.

60. Lee W, Nims RJ, Savadipour A, et al. Inflammatory signaling sensitizes Piezo1 mechanotransduction in articular chondrocytes as a pathogenic feed-forward mechanism in osteoarthritis. Proc Natl Acad Sci U S A. 2021;118(13).

61. Brylka LJ, Alimy AR, Tschaffon-Muller MEA, et al. Piezo1 expression in chondrocytes controls endochondral ossification and osteoarthritis development. Bone Res. 2024;12(1):12.

62. Coste B, Houge G, Murray MF, et al. Gain-of-function mutations in the mechanically activated ion channel PIEZO2 cause a subtype of Distal Arthrogryposis. Proc Natl Acad Sci U S A. 2013;110(12):4667–4672.

63. Ma Y, Zhao Y, Cai Z, Hao X. Mutations in PIEZO2 contribute to Gordon syndrome, Marden-Walker syndrome and distal arthrogryposis: A bioinformatics analysis of mechanisms. Exp Ther Med. 2019;17(5):3518–3524.

64. Dombroski JA, Hope JM, Sarna NS, King MR. Channeling the Force: Piezo1 Mechanotransduction in Cancer Metastasis. Cells. 2021;10(11).

65. Wang Q, Yu Y, Liang X, Wan D, Du K, Zhu P. Pan-cancer analysis of PIEZO1: a promising biomarker for diagnosis, prognosis, and targeted therapies. Front Immunol. 2025;16:1625734.

66. Szczot M, Liljencrantz J, Ghitani N, et al. PIEZO2 mediates injury-induced tactile pain in mice and humans. Sci Transl Med. 2018;10(462).

67. Wan Y, Zhou J, Li H. The Role of Mechanosensitive Piezo Channels in Chronic Pain. J Pain Res. 2024;17:4199–4212.

68. Obeidat AM, Wood MJ, Adamczyk NS, et al. Piezo2 expressing nociceptors mediate mechanical sensitization in experimental osteoarthritis. Nat Commun. 2023;14(1):2479.

69. Behunova J, Gerykova Bujalkova M, Gras G, et al. Distal Arthrogryposis with Impaired Proprioception and Touch: Description of an Early Phenotype in a Boy with Compound Heterozygosity of PIEZO2 Mutations and Review of the Literature. Mol Syndromol. 2019;9(6):287–294.

70. Wairimu NW, Wairagu P, Chepukosi KW, et al. Sodium Metabisulfite-Induced Hematotoxicity, Oxidative Stress, and Organ Damage Ameliorated by Standardized Ginkgo biloba in Mice. J Toxicol. 2023;2023:7058016.

