## Supplementary file for "Cannabidiol Inhibits PIEZO Channels to Mitigate Red Blood Disorders"

*Correspondence:

Huanghe Yang,

**The file includes:**

Supplementary Methods
Supplementary Figure 1 to Figure 6
Captions for Supplementary Video 1 to Video 4

### SUPPLEMENTARY METHODS

#### Cell lines

HEK293T cells were cultured in high-glucose DMEM medium (Gibco, REF 11995-065), supplemented with 10% FBS and 1% penicillin/streptomycin. 100 μg/mL hygromycin (Corning, Cat. #30-240-CR) was added to the culture medium to maintain stable expression of mTMEM16F in the stable HEK293T cell line. HEK293T-P1KO cells (Piezo1 knockout human embryonic kidney cells) were obtained from Ardem Patapoutian[1]. All cells were cultured in a humidified atmosphere with 5% $CO_2$ at 37°C.

Human umbilical vein endothelial cells (HUVECs) were cultured in EGM-2 BulletKit (Lonza, #CC-3162, authenticated by the Duke Cell Culture Facility) in a humidified incubator at 37°C and 5% $CO_2$-95% air. For the adhesion assay, cells were seeded in ibidi-treat μ-Slide VI 0.4 (Cat.No: 80606) following the manufacturer's instructions. Briefly, 30 μl of $8\times10^5$ cells/ml HUVECs were seeded in each channel. After cell attachment, each reservoir was filled with 60 μl medium. The adhesion assay was performed after a confluent monolayer was formed.

#### Molecular biology

The human PIEZO1-IRES-eGFP construct was a generous gift from Dr. Philip A. Gottlieb (University at Buffalo, SUNY). The R2488Q mutation was generated by PCR-based site-directed mutagenesis. Cells were transfected with XtremeGENE 9 DNA Transfection Reagent (Roche, #06365787001) or Lipofectamine 2000 (ThermoFisher, #11668019) according to the manufacturer's instructions. For a 24-well plate, 500 ng of plasmid DNA encoding human wild-type PIEZO1, mutant PIEZO1, or wild-type PIEZO2 was used per well.

#### Flow cytometry

RBCs were centrifuged at 100 × *g* for 12 min, and the pellets were washed twice with PBS to obtain packed RBCs. Packed cells were then diluted 1:20 with Hank's balanced salt solution (HBSS). A 3 μL aliquot of the diluted RBC suspension was added to 200 μL HBSS containing CF488-conjugated Annexin V (1:125). Designed concentrations of Yoda1 were added, and samples were incubated for 10 min at room temperature. Samples were analyzed using a BD FACS Canto flow cytometer (BD Biosciences, Franklin Lakes, NJ) following the manufacturer's guidelines. Erythrocytes were identified by gating on forward and side scatter profiles. Excitation was performed with a 488 nm laser, and green fluorescence emission was collected through a 530 nm filter. Results were expressed either as the percentage of Annexin V-positive erythrocytes or as the mean fluorescence intensity (MFI) of Annexin V staining. For pharmacological testing, RBCs were pre-incubated with drugs of interest prior to Yoda1 addition, followed by a 10 min incubation at room temperature. Flow cytometry data were analyzed using FCS Express software (De Novo Software, Pasadena, CA).

**Fluorescence imaging of $Ca^{2+}$ and PS exposure**

TMEM16F-stable HEK293T cells were seeded on poly-L-lysine-coated coverslips (MilliporeSigma) and maintained in DMEM supplemented with hygromycin B (100 μg/mL; Corning, REF 30-240-CR) to preserve TMEM16F expression. For PIEZO1 expression, cells were transiently transfected with human PIEZO1-IRES-eGFP (500 ng per well, 24-well plate) using XtremeGENE 9 (Roche, REF 06365787001) according to the manufacturer's instructions, and cultured for 24 h before imaging.

For simultaneous monitoring of intracellular $Ca^{2+}$ and phosphatidylserine (PS) exposure, cells on coverslips were loaded with Calbryte 590 AM (1 μM in HBSS) for 15 min at 37 °C, 5% $CO_2$. Coverslips were then transferred to imaging buffer containing Annexin V conjugate (1:140

dilution) on glass slides. Cells exhibiting both membrane and cytosolic GFP fluorescence were recorded for a 50-s baseline, followed by application of Yoda1 (2 µM) to activate PIEZO1. A Zeiss LSM 780 inverted confocal microscope was used to acquire morphology, $Ca^{2+}$ dynamics, and PS exposure in real time at 5-s intervals. RBCs were placed on coverslips in buffer (140 mM NaCl, 5 mM KCl, 2 mM $MgCl_2$, 10 mM HEPES, 2 mM $CaCl_2$, pH 7.4) and loaded with Calbryte 590 AM (1 µM; AAT Bioquest, cat. 20701) for 30 min at 37 °C, 5% $CO_2$. Coverslips were then transferred to buffer containing Annexin V CF488A (1:140; Biotium, cat. 29005). After a 50-s baseline, Yoda1 (2 µM for human RBCs) was added to stimulate PIEZO1. Imaging was performed on a Zeiss LSM 780 inverted confocal microscope with 5-s sampling intervals to capture morphology, $Ca^{2+}$ transients, and PS exposure. MATLAB code for quantifying CaPLSase activity from fluorescence imaging [2-5] is available at: https://github.com/yanghuanghe/scrambling_activity.

**Intracellular $Ca^{2+}$ measurement using a plate reader**

Intracellular $Ca^{2+}$ levels were quantified using the ratiometric dye Fura-2 AM. Briefly, 2 µM Fura-2 AM was added to either 100 µL of packed RBCs diluted in 900 µL HBSS. Samples were incubated at 37 °C for 45 min to allow dye loading. Cells were then centrifuged at 200 × *g* for 5 min to remove unbound dye, resuspended in fresh HBSS, and incubated for an additional 10-15 min at 37 °C to allow de-esterification. Fura-2 fluorescence was measured using a SpectraMax M5 plate reader (Molecular Devices) with dual excitation at 340 nm and 380 nm, and emission detection at 510 nm. Baseline fluorescence was recorded for 1 min before the addition of Yoda1 at the indicated concentration. Fluorescence readings were acquired every 30 s for 10-20 min, and data were analyzed using Microsoft Excel.

**Yoda1-induced hemolysis assay**

Packed RBCs (20 μL) were added to 1 mL HBSS ± CBD. Yoda1 (final 2 μM) was then introduced, and samples were incubated at room temperature. At designated time points, 100 μL of clarified supernatant was transferred to a 96-well plate. At the end of the experiment (~2 h), absorbance was measured at 577 nm with 690 nm for background subtraction. For the positive control (complete lysis), 20 μL packed RBCs from each sample were added to 1 mL distilled $H_2O$ for 5 min, then the supernatant was collected after brief centrifugation. Samples without Yoda1 served as the negative control.

**Centrifugation-induced dehydration assay**

Packed RBCs (20 μL) were added to 1 mL HBSS with or without test compounds and spun at 900 × g for 12 min. For each condition, bright-field images were captured from three random fields and averaged as a single data point after morphology analysis.

**RBC-endothelium adhesion assay**

PBS was pre-warmed to 37 °C. Yoda1 was prepared at 2 μM in 1 mL HBSS. Packed RBCs (10 μL) were diluted 1:100 in 1 mL HBSS and labeled with Calcein AM (1 μM, 15 min, 37 °C). Labeled cells were centrifuged (200 × g, 5 min) to remove excess dye. One microliter of packed RBCs was then resuspended in the 2 μM Yoda1 solution to achieve a 1:1000 final dilution and incubated for 10 min at room temperature. Cells were pelleted again (200 × g, 5 min) to remove free Yoda1 and resuspended in 1 mL pre-warmed PBS. For flow experiments (ibidi, #80606), air bubbles were cleared by briefly running a syringe pump (Chemyx Fusion 200) at ~5 mL $min^{-1}$. Approximately 0.5 mL of the RBC suspension was introduced into the chamber and incubated for 10 min at 37 °C to allow adhesion. Non-adherent cells were removed by perfusion at 1 dyne $cm^{-2}$ for 5 min. Adherent RBCs were imaged immediately on an Olympus IX83 microscope. For quantification, three representative fields per condition were analyzed in ImageJ, and the mean number of adherent RBCs across these fields was used for statistical analysis.

**Prothrombinase Assay**

Packed RBCs (1 µL) were resuspended in 500 µL assay buffer containing 3 mM $Ca^{2+}$ and incubated with 2 µM Yoda1 (from 1 mM stock; 1 µL in 500 µL) for 10 min at room temperature. Factor Xa (1 nM) and Factor Va (2 nM) were added (from 100 nM and 200 nM stocks; 5 µL each into 500 µL) and incubated for 2 min at 37 °C. Then, 50 µL of the reaction was transferred to black 96-well plates, and prothrombin (1.4 µM) was added (from 36 µM stock; 2 µL/well). After 2 min at 37 °C, 100 µL quench buffer was added to stop the reaction. The fluorogenic thrombin substrate Z-Gly-Gly-Arg-AMC (Bachem) was prepared at 80 mM in DMSO and added to a final concentration of 5 µM (0.75 µL/well). Fluorescence (Ex 355 nm/Em 460 nm) was recorded every 15-30 s, with plate shaking before and between reads to ensure mixing.

**Electrophysiology**

Ionic currents were recorded using an Axopatch 200B amplifier (Molecular Devices) and acquired with the *pClamp* software suite (Molecular Devices) in cell-attached, outside-out, or inside-out configurations, as indicated. Patch pipettes were fabricated from borosilicate glass capillaries (Sutter Instruments) and fire-polished with a microforge (Narishige) to a resistance of 2-3 MΩ. For current-voltage (I-V) relationships, currents were evoked by a voltage-step protocol from −100 mV to +160 mV in 20 mV increments, with a holding potential of −60 mV.

The standard bath solution contained (in mM): 140 CsCl, 10 HEPES, and 0 or 2.5 $CaCl_2$, as indicated, adjusted to pH 7.4 with CsOH. For cell-attached and inside-out recordings, the pipette solution contained (in mM): 140 CsCl, 10 HEPES, and 1 $MgCl_2$ (pH 7.4, CsOH), while the bath and perfusion solutions contained 140 CsCl and 10 HEPES (pH 7.2, CsOH). Mechanical stimulation was applied using a high-speed pressure-clamp system (ALA Scientific Instruments, model HSPC-2-SB), and patches were held at −80 mV.

For outside-out recordings of TMEM16F currents, the pipette solution contained (in mM): 140 CsCl, 10 HEPES, 1 $MgCl_2$, and 0.1 free $Ca^{2+}$ (pH 7.2, CsOH), while the bath solution consisted of 140 CsCl and 10 HEPES (pH 7.2, CsOH). All recordings were performed at room temperature (22-25 °C).

**Quantification of I-V relation and dose-response to compounds**

*I-V* curves were constructed from the steady-state peak currents. Individual *I-V* curves were fitted with a Boltzmann function,

$$I(V) = \frac{I_{max}}{1+e^{\frac{-ZF(V-V_{0.5})}{RT}}} \qquad (2)$$

where $I_{max}$ denotes the fitted value for maximal conductance at a given voltage, $V_{0.5}$ denotes the voltage of half maximal activation of conductance, $z$ denotes the net charge moved across the membrane during the transition from the closed to the open state and $F$ denotes the faraday constant.

Dose response curves were fitted with Hill equation,

$$\frac{F}{F_{max}} = \frac{1}{1+\frac{[IC_{50}]}{[drug]}^{H}} \qquad (3)$$

where $F/F_{max}$ denotes the corresponding signal normalized to the max fluorescence signal at given stimuli, [drug] denotes free drug concentrations, $H$ denotes Hill coefficient, and $IC_{50}$ denotes the half-maximal inhibition concentration of the individual drug.

**SUPPLEMENTARY FIGURES**

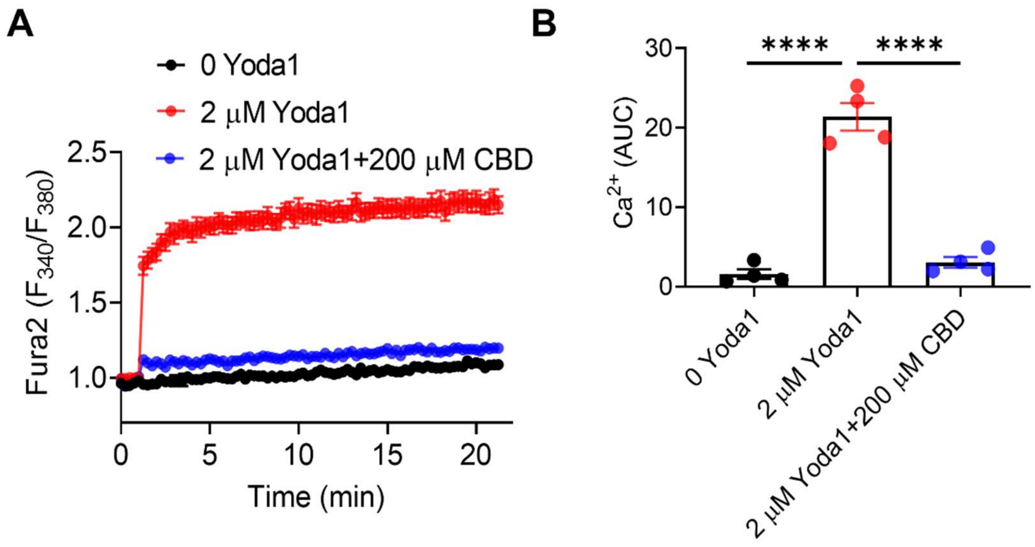


**Figure S1. 200 μM CBD suppresses Yoda1-induced $Ca^{2+}$ increase in human RBCs.** (A) Fura-2 measurements of Yoda1-induced $Ca^{2+}$ influx in healthy donor (HD) RBCs with or without CBD (200 μM). RBCs without Yoda1 served as a negative control. Data are shown as mean ± SEM from three replicates. (B) Quantification of total $Ca^{2+}$ influx from panel A, expressed as area under the curve (AUC). Each dot represents the average of three replicates per blood sample (N = 4 donors). Statistical analysis: unpaired two-sided Student's *t* test; ****P < .0001.

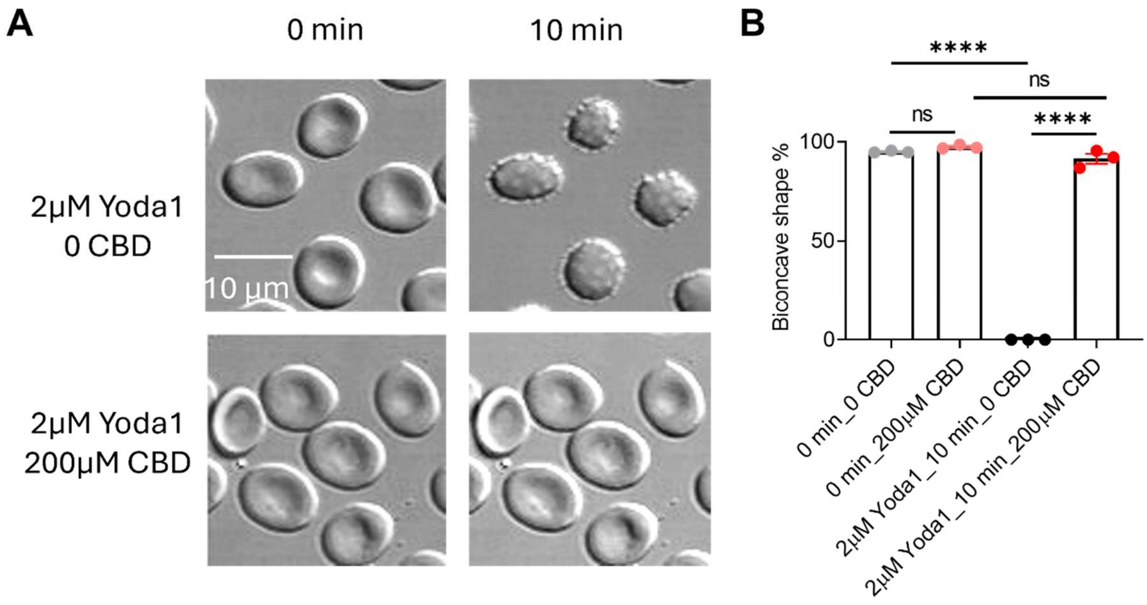


**Figure S2. 200 µM CBD prevents Yoda1-induced RBC dehydration.** (A) Representative images of morphological changes in RBCs treated with Yoda1 (2 µM) in the absence or presence of CBD (200 µM), captured at 0 and 10 minutes after Yoda1 treatment. (B) Quantification of biconcave-shaped RBCs with or without CBD at 0 and 10 minutes after Yoda1 stimulation. Statistical analysis: one-way ANOVA followed by Tukey's test; ****P < 0.0001; ns, not significant (N = 3 biological replicates).

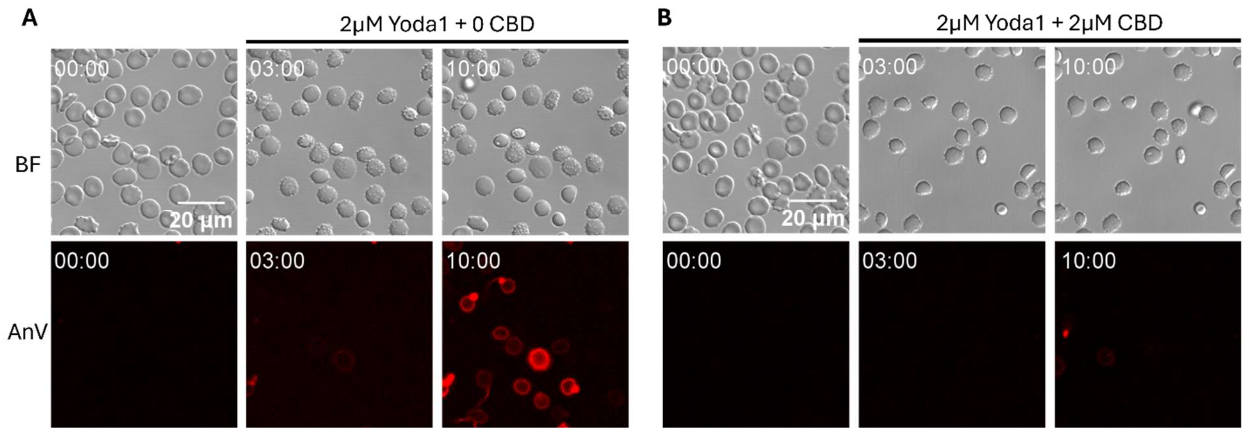


**Figure S3. 2 µM CBD suppresses Yoda1-induced PS exposure without affecting Yoda1-induced morphological changes.** (A-B) Representative images of morphological changes (Bright field/BF, top) and PS exposure (AnV, bottom) in RBCs treated with 2 µM Yoda1 in the absence (A) or presence of 2 µM CBD (B), captured at 0, 3 and 10 minutes after Yoda1 treatment.

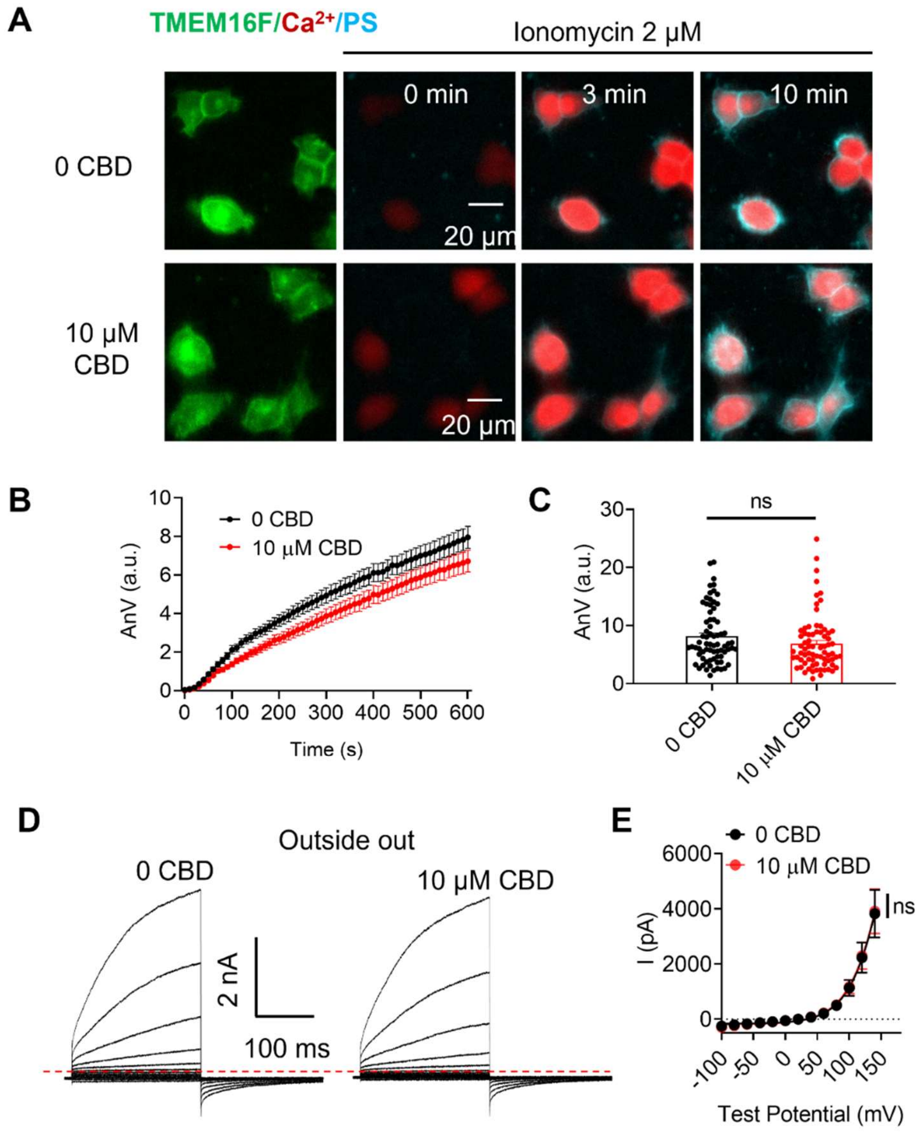


**Figure S4. 10 µM CBD does not affect TMEM16F activity in HEK293T cells.** (A) Representative images of TMEM16F-mediated PS exposure in HEK293T cells stably expressing TMEM16F, with or without extracellular CBD (10 µM). Images were acquired every 10 seconds following ionomycin (2 µM) treatment. (B) Time course of TMEM16F-mediated PS exposure in the presence and absence of CBD. Data are presented as mean ± SEM (n = 72 and 73 cells for 0 and 10 µM CBD, respectively; 3 biological replicates). (C) Comparison of AnV intensity at 10 minutes with and without CBD. Statistical analysis: unpaired two-sided Student's *t* test; ns, not significant (n = 72 and 73 cells for 0 and 10 μM CBD, respectively; 3 biological replicates). (D) Outside-out patch clamp recording of TMEM16F current elicited by 0.1 mM $Ca^{2+}$ in the pipette solution with and without extracellular CBD (10 μM). (E) The I-V relationship of TMEM16F current in D. Statistical analysis: unpaired two-sided Student's *t* test; ns, not significant (n = 6 cells for both 0 and 10 μM CBD groups).

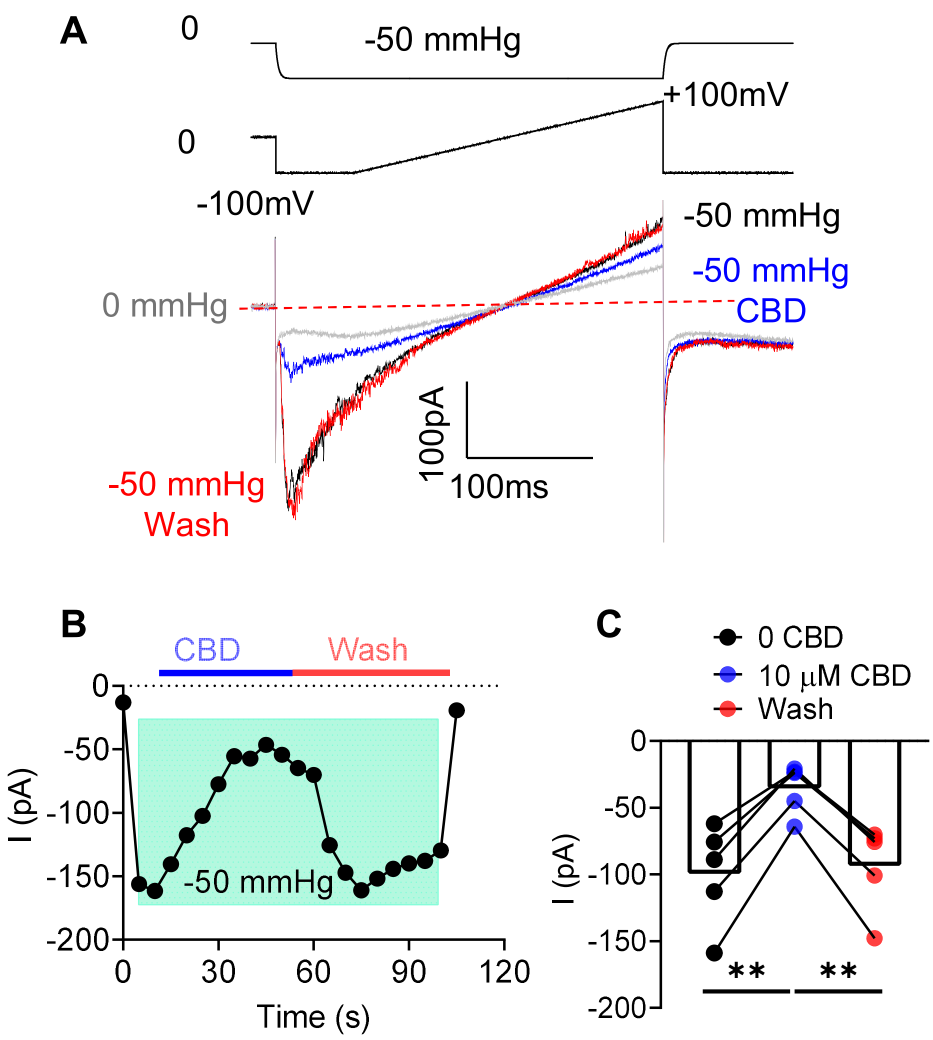


**Figure S5. CBD can block PIEZO1 from the intracellular side.** (A) Representative current traces showing the effect of CBD on PIEZO1 activity. Currents were elicited using a voltage ramp protocol (-100 mV to +100 mV) from a holding potential of 0 mV, under -50 mmHg negative pressure. The gray trace represents baseline at 0 mmHg; the black trace, current at -50 mmHg; the blue trace, current in the presence of 10 µM CBD; and the red trace, current after CBD washout. (B) Time course of CBD-mediated inhibition of PIEZO1 currents and recovery upon washout. A -50 mmHg pressure pulse was applied during the shaded interval.

(C) Quantification of PIEZO1 currents at -50 mmHg under the indicated conditions. Data are presented as mean ± SEM (N = 5 biological replicates). Statistical analysis was performed using one-way ANOVA followed by Tukey's post hoc test (**$P < 0.01$; ns, not significant).

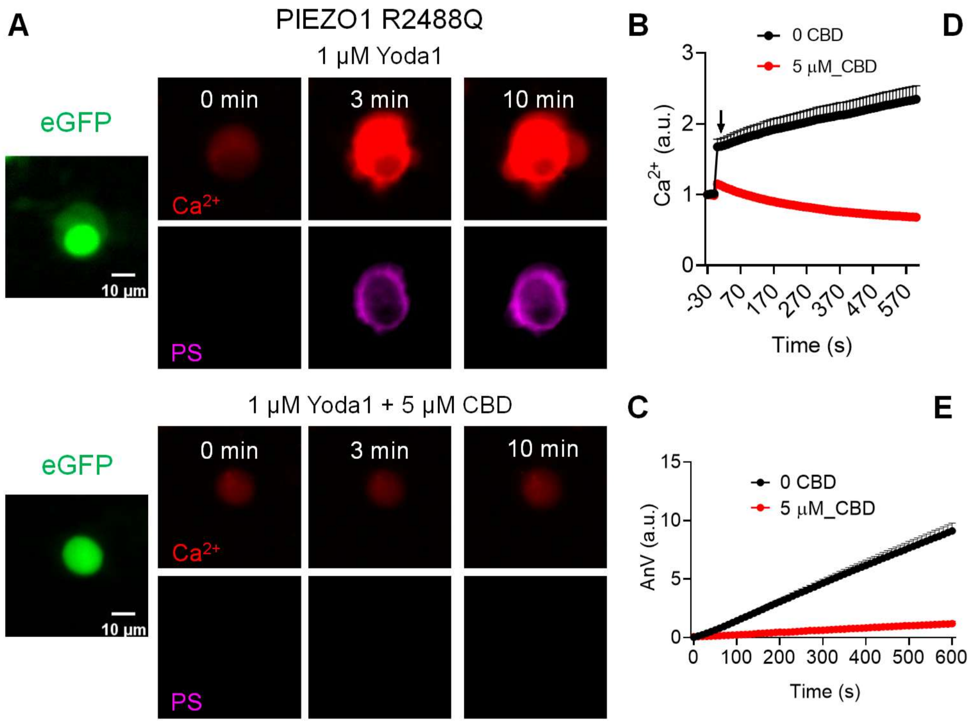


**Figure S6. CBD inhibits PIEZO1 gain-of-function mutation-induced $Ca^{2+}$ influx and phosphatidylserine exposure.** (A) Representative images of Yoda1 (1 μM)-induced $Ca^{2+}$ dynamics (red) and PS exposure monitored by Annexin V (AnV, magenta) in WT HEK293T cells overexpressing the PIEZO1-R2488Q HX mutation (green) without CBD (top) and with 5 μM CBD (bottom). (B–C) Time-dependent Yoda1-stimulated $Ca^{2+}$ signal (B) and AnV-reported lipid scrambling (C) in the absence or presence of CBD. Data are mean ± SEM (n = 107 and 154 cells, from ≥3 biological replicates, without and with CBD, respectively). (D–E) Quantification of $Ca^{2+}$ influx (D) and lipid scrambling (E) with and without CBD (5 μM). Data are mean ± SEM. Statistical significance was determined by unpaired two-sided Student's t test (****P < 0.0001; n = 107 and 154 cells from ≥3 biological replicates, without and with CBD, respectively).

### SUPPLEMENTARY VIDEOS

**Supplementary Video 1.** This video shows simultaneous imaging of RBC morphology, $Ca^{2+}$ (Calbryte 520, green) and fluorescently tagged Annexin V (AnV-CF594, red) in health donor (HD) RBCs upon Yoda1 stimulation (2 μM). Images were acquired every 5 seconds for ~10 minutes. Yoda1 was added at time 0.

**Supplementary Video 2.** This video shows simultaneous imaging of RBC morphology, $Ca^{2+}$ (Calbryte 520, green) and fluorescently tagged Annexin V (AnV-CF594, red) in health donor (HD) RBCs upon Yoda1 stimulation (2 μM) in the presence of CBD (200 μM). Images were acquired every 5 seconds for ~10 minutes. Yoda1 was added at time 0.

**Supplementary Video 3.** Tape response assay (see Methods for details) performed on control mice following intraperitoneal injection of saline. The original 2-minute recording is shown at 4× speed.

**Supplementary Video 4.** Tape response assay (see Methods for details) performed on mice after intraperitoneal injection of 10 mg/kg CBD. The original 2-minute recording is shown at 4× speed.

**References:**


1. Dubin AE, Murthy S, Lewis AH, et al. Endogenous Piezo1 Can Confound Mechanically Activated Channel Identification and Characterization. *Neuron*. 2017;94(2):266-270.e263.
2. Le T, Jia Z, Le SC, Zhang Y, Chen J, Yang H. An inner activation gate controls TMEM16F phospholipid scrambling. *Nature Communications*. 2019;10:1846.
3. Zhang Y, Le T, Grabau R, et al. TMEM16F phospholipid scramblase mediates trophoblast fusion and placental development. *Science Advance*. 2020;6.
4. Liang P, Yang H. Molecular underpinning of intracellular pH regulation on TMEM16F. *J Gen Physiol*. 2021;153(2).
5. Liang P, Wan Y-CS, Shan KZ, et al. Targeting PIEZO1-TMEM16F Coupling to Mitigate Sickle Cell Disease Complications. *American Journal of Hematology*. 2025;n/a(n/a).
